# Direct inference of the distribution of fitness effects of spontaneous mutations in *Chlamydomonas reinhardtii*

**DOI:** 10.1101/571018

**Authors:** Katharina B. Böndel, Susanne A. Kraemer, Tobias S. Samuels, Deirdre McClean, Josianne Lachapelle, Rob W. Ness, Nick Colegrave, Peter D. Keightley

## Abstract

Spontaneous mutations are the source of new genetic variation and are thus central to the evolutionary process. In molecular evolution and quantitative genetics, the nature of genetic variation depends critically on the distribution of fitness effects (DFE) of mutations. Spontaneous mutation accumulation (MA) experiments have been the principal approach for investigating the overall rate of occurrence and cumulative effect of mutations, but have not allowed the effects of individual mutations to be studied directly. Here, we crossed MA lines of the green alga *Chlamydomonas reinhardtii* with its unmutated ancestral strain to create haploid recombinant lines, each carrying an average of 50% of the accumulated mutations in a variety of combinations. With the aid of the genome sequences of the MA lines, we inferred the genotypes of the mutations, assayed their growth rate as a measure of fitness, and inferred the DFE using a novel Bayesian mixture model that allows the effects of individual mutations to be estimated. We infer that the DFE is highly leptokurtic (L-shaped), and that a high proportion of mutations increase fitness in the laboratory environment. The inferred distribution of effects for deleterious mutations is consistent with a strong role for nearly neutral evolution. Specifically, such a distribution predicts that nucleotide variation and genetic variation for quantitative traits will be insensitive to change in the effective population size.

## Introduction

Understanding evolution requires an understanding of the origin of new genetic variation from mutation. This includes knowing the rates of mutation at individual loci and the magnitudes of their effects on fitness and other traits. Of particular interest is the distribution of fitness effects for new mutations (the DFE), describing the relative rates of occurrence of mutations with different selective effect sizes. The DFE informs about the frequencies of small-*versus* large-effect mutations and the frequencies of advantageous *versus* deleterious mutations, and is therefore of fundamental importance in population and quantitative genetics. For example, the DFE appears in the nearly neutral model of molecular evolution (Ohta 1977), where deleterious mutations are effectively selected against in large populations, but behave as selectively neutral in small populations. Kimura (1979; 1983) showed that if the DFE is strongly leptokurtic (L-shaped) molecular genetic variation at sites subject to natural selection increases slowly with increasing effective population size (*N_e_*), and molecular evolution is potentially constant between species with different effective size. This is therefore broadly consistent with empirical observations. The DFE is also important for predicting selection response for quantitative traits and the nature of quantitative genetic variation (Robertson 1967). For example, the contribution of mutation to response to selection depends critically on the shape of the DFE, the response occurring more quickly with more leptokurtic distributions (Hill 1982). Analogously with the relationship between nucleotide variation and *N_e_*, genetic variation for fitness (or a trait correlated with fitness) is predicted to increase slowly as a function of *N*_*e*_ if the DFE is leptokurtic (Keightley and Hill 1990), and could thus explain why genetic variation for quantitative traits is apparently relatively invariant between species (Potsma 2014).

In the light of its fundamental importance, there has been much previous work aimed at inferring the DFE. Two different approaches have principally been applied for spontaneous mutations occurring in the whole genome (rather than just a single locus): the analysis of nucleotide polymorphism data from a sample of individuals from a population, and spontaneous mutation accumulation (MA) experiments (reviewed by Eyre-Walker and Keightley 2007). Under the former approach (Eyre-Walker et al 2006; Keightley and Eyre-Walker 2007; Boyko et al. 2008; Tataru et al 2017), the site frequency spectra for putatively neutral and selected sites (typically synonymous and nonsynonymous sites of protein-coding genes, respectively) are compared, and parameters of the DFE for the mutations at the selected sites inferred. The approach makes several assumptions, notably that variation at the selected sites is explained by a balance between an input of new deleterious mutations, natural selection and genetic drift, and that selection is absent from the putatively neutral class of sites. It is only capable of inferring the DFE for mutations that stand an appreciable chance of segregating in the sample of individuals from the population, implying that inferences are only relevant to mutations with effects that are not substantially greater than 1/*N_e_*. This can be an extremely small value if *N*_*e*_ is large. Furthermore it can only be applied to specific functional categories of sites in the genome.

In a spontaneous MA experiment, sublines of the same initial genotype are maintained at small effective population size in the near absence of natural selection for many generations, allowing mutations to accumulate effectively at random. The DFE can be estimated using the among-MA line distribution of phenotypic values for traits related to fitness (such as fecundity or viability) (Keightley 1994; García-Dorado 1997; Shaw et al 2002). The information that can be obtained by this approach is extremely limited, however, principally because the numbers of mutations carried by individual lines are not included in the analysis, so an overall genomic rate parameter has to be estimated, and this is highly confounded with the DFE parameters (Keightley 1998; Halligan and Keightley 2009).

Genome sequencing technology now allows the identification of the nearly complete complement of mutations carried by a set of MA lines, and in combination with phenotypic information this can potentially be used to leverage information on the DFE (Katju and Bergthorsson 2018). Previous analysis of spontaneous MA experiments have, however, only studied the cumulative effects of new mutations, whereas accurate inference requires estimation of the effects of individual mutations. For example, we have shown that there is a negative correlation between the number of new mutations carried by a MA line and fitness, but this gives only limited information on the DFE (Kraemer et al. 2017).

Previously, we carried out a spontaneous MA experiment in the single-celled green alga *Chlamydomonas reinhardtii* for ~1,000 generations, have measured fitness-related traits in a range of environmental conditions (Morgan et al 2014; Kraemer et al 2016; 2017), and have employed genome sequencing to determine the complement of mutations carried by the lines (Ness et al 2015). Here, we have crossed six of these *C*. *reinhardtii* MA lines of the CC-2931 genetic background with a compatible ancestor of the same background genotype, but of the opposite mating type. We thereby generated 1,526 recombinant lines (RLs), each carrying an average of 50% of the mutations of the MA line parent in different combinations. We genotyped the RLs at the locations of the known mutations and assayed their growth rate as a measure for fitness. Across the six lines there are nearly 400 unique mutations, so an analysis where each mutation is treated as a fixed effect is not appropriate. Instead, we developed a MCMC approach with a random effects model in which mutation effects are assumed to be sampled from some distribution or a mixture of distributions. We investigate a number of distributions to infer the distribution of effects for the individual mutations on growth rate. We show that the DFE is highly leptokurtic (L-shaped) and that a high proportion of mutations increase fitness in the laboratory environment.

## Results

To directly infer the DFE, we crossed six *C. reinhardtii* MA lines derived from the CC-2931 strain to an ancestral strain of the same genetic background and the opposite mating type to produce a total of 1,526 recombinant lines (RLs) (Table 1, Table S1). We genotyped 386 of the 476 mutations detected in our previous whole-genome sequencing study (Ness et al. 2015) (Table 1, Table S2). Among the 681 different recovered haplotypes, mutations were present at an average frequency of 49.1% (10.3% - 85.4%), which is close to the expected average of 50% (Figure S1). The number of haplotypes obtained for each MA line and their frequencies were quite variable, however (Table S3). For example, we obtained 214 haplotypes for MA line L03, and no haplotype was found more than four times, whereas we obtained only 67 haplotypes for MA line 14 and one of these haplotypes was found 18 times.

**Table 1.** Data overview.

| *MA line cross* | *Number of Mutations* | *Number of RLs* | *Number of haplotypes* |
| --- | --- | --- | --- |
| L03 | 39 | 247 | 214 |
| L06 | 69 | 238 | 109 |
| L07 | 59 | 261 | 69 |
| L09 | 98 | 272 | 68 |
| L11 | 66 | 272 | 154 |
| L14 | 55 | 236 | 67 |
| Combined | 386 | 1,526 | 681 |

### Relationship between number of mutations and growth rate

As a measure of fitness, we assayed the maximum growth rate of each RL, the parental MA lines and the unmutated ancestral strain in liquid culture. To determine whether mutations have an overall directional effect on fitness, we used mixed models to test for a relationship between the number of mutations carried by RLs and their ancestors and fitness (Figure 1). In the case of only one of the six MA lines (L03), including number of mutations leads to a significantly better fit (*P* = 0.0008; Table 2), and the improvement in fit for an analysis of the combined data set of all six MA line crosses is nonsignificant (*P* = 0.080). This could either mean that there is insufficient power to detect mutational effects, or that there is a mixture of mutations with positive and negative effects on fitness. The latter explanation is supported, because there is a highly significant between-haplotype component of variation for the trait (*P* < 2.2×10^−16^ for the whole data set; *P* between 4.1×10^−13^ (L14) and 0.022 (L06) for the individual MA lines). We repeated this analysis fitting number of mutations of specific types (SNP, indel, exonic, intronic, intergenic; Table S4). Including the number of mutations gave a significantly better fit in the cases of MA line L03 for all mutation types except intronic, for MA line L11 for intronic mutations and for the whole data for exonic mutations.

**Table 2.**
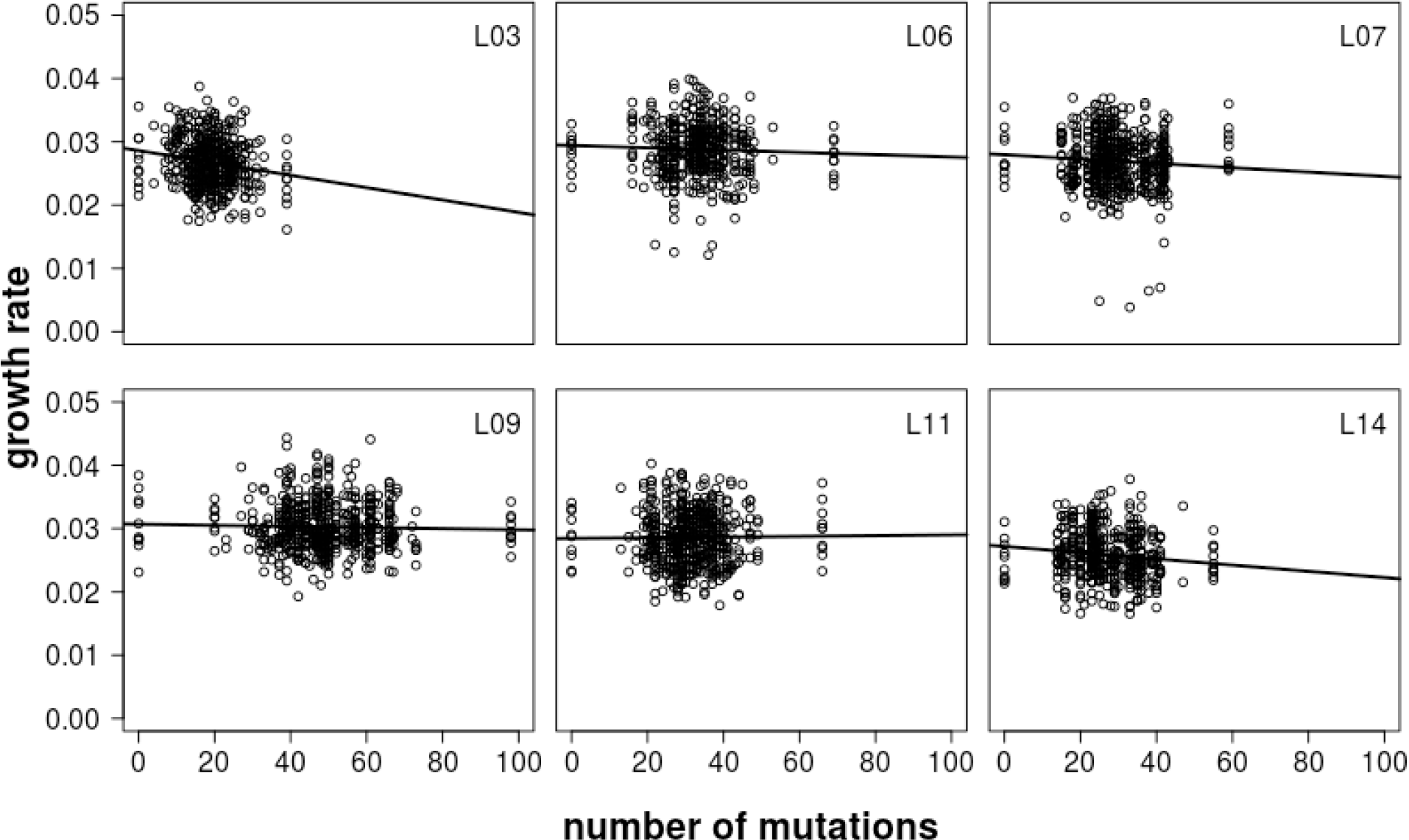
Likelihood ratio tests for mixed model analysis of growth rate as a function of number of mutations of all kinds with 1 degree of freedom.

**Figure 1.**
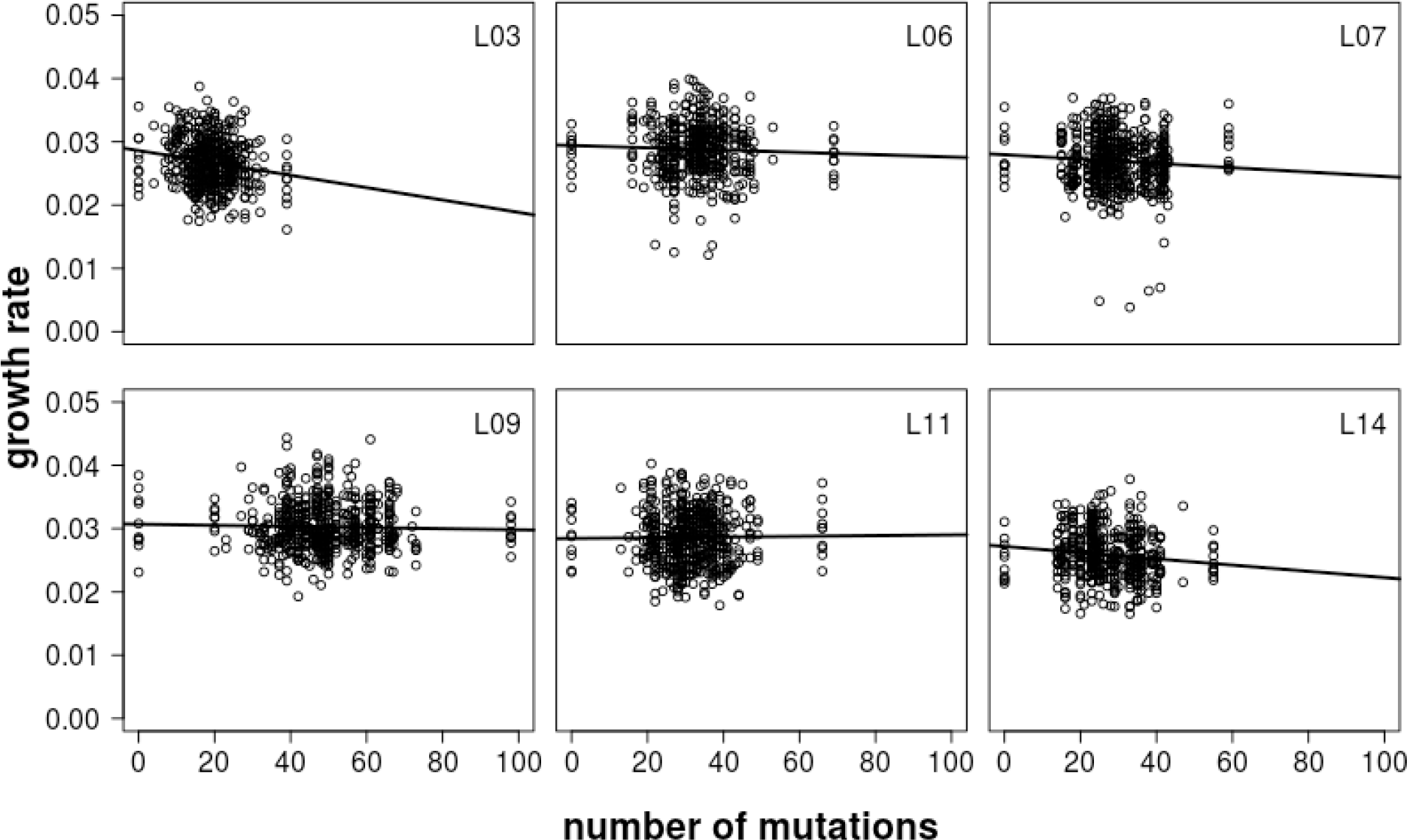
Relationship between growth rate and number of mutations carried by a RL or ancestor for the six CC-2931 MA line crosses. Linear regression lines are shown.

### Inference of the DFE by MCMC assuming mutation effects fall into discrete categories

The relationship between mutation number and fitness tells us little about the DFE for individual mutations. We therefore developed an approach that allows the DFE be estimated in a Bayesian mixture model (implemented by Markov Chain Monte Carlo, MCMC). This assumes that the effects of mutations either come from a mixture of point masses or a mixture of gamma distributions. To maximize power, we focussed much of the analysis on a merged data set of all six MA line backcrosses.

We first examined whether there is evidence for an overall directional effect of new mutations on growth rate by running the analysis while assuming a two category model with one non-zero effect category (effect = *e*_1_, proportion = *q*_1_) and one zero-effect category (i.e., *e*_0_ = 0, *q*_0_ = 1 - *q*_1_). The results (Table 3, Figure S2, S3) suggest that there is an appreciable frequency (~4%) of mutations reducing growth rate by ~3%, whereas the majority of mutations are allocated to the zero-effect category.

**Table 3.** Bayesian MCMC estimates and 95% credible intervals for mutation effect (*e*) and mutation frequency (*q*) parameters under two or three-category models. Both models include a class of mutations with zero effect on the trait.

| *Parameter* | *Mutation categories in model* | *Estimate* | *95% credible interval* |  |
| --- | --- | --- | --- | --- |
| $e_1$ | 2 | -0.031 | -0.044 | -0.023 |
| $q_1$ | 2 | 0.042 | 0.020 | 0.079 |
| $e_1$ | 3 | -0.024 | -0.043 | -0.011 |
| $q_1$ | 3 | 0.071 | 0.031 | 0.42 |
| $e_2$ | 3 | 0.021 | 0.010 | 0.068 |
| $q_2$ | 3 | 0.048 | 0.010 | 0.41 |

We then analysed the combined data set assuming a model with three categories of mutational effects (one zero-effect category and two finite effect categories, *e*_1_ and *e*_2_). As expected, given that the two-category model supports the presence of negative mutational effects, a category of negative effects is inferred (*e*_1_, Table 3; Figure S4, S5). This has a similar posterior mode as the two-category model, but the credible interval is somewhat wider, as expected for a more data rich model. There is also support for a class of positive-effect mutations (*e*_2_, Table 3), which has a somewhat lower absolute modal estimate than the negative category. The estimated frequency of positive-effect mutations is only slightly lower than that of the negative-effect mutations. Results from the analysis of data from individual MA line crosses (Table S5) are consistent with the presence of a mixture of negative- and positive-effect mutations.

We then analysed data sets in which phenotypic values were permuted within plate. As expected under the null model, the distributions of estimated values of *e*_1_ and *e*_2_ centre on zero and the estimates of *e*_1_ and *e*_2_ from the real data are well outside the distributions obtained from permuted data (Figure S6).

Analysis of a model with four categories of effects (one of which is a zero-effect category) also gives negative and positive posterior modes for two classes of mutational effects *e*_1_ and *e*_2_. However, it is difficult to determine whether there is an additional mutational class *e*_*3*_ that is different from the zero-effect class or *e*_1_ and *e*_2_ because of the presence of label switching (Jasra et al 2005) between the three classes of effects and their frequencies (data not shown).

### Two-sided gamma DFE model

Although informative about the overall directional effects of mutations, models in which mutations fall into discrete categories are unrealistic, because they assume no variance among the effects of mutations within each category. We therefore analyzed the combined data set for the six MA line crosses under a two-sided gamma distribution of effects, which assumes that the effects of mutations are continuously distributed. We assumed the gamma distribution, because it is a flexible two-parameter distribution (*α* = scale, *β* = shape) that can take a wide variety of shapes, ranging from a highly leptokurtic, L-shaped distribution (*β* → 0) to a point mass (*β* → ∞). We assumed that positive- and negative-effect mutations can have different absolute means, but their distributions have the same shape parameter. The results from the analysis of the combined data set (Table 4; Figure S7) suggest that the DFE is highly leptokurtic (i.e., *β* is close to 0.3), and that the means for positive-and negative-effect mutations (= *β*/*α*) are very small, reflecting the concentration of mutations with effects close to zero. Consistent with the analysis assuming discrete classes of mutations (Table 3), there is a substantial proportion of positive-effect mutations (i.e., ~80%; Table 4). The estimated DFE for the two-sided gamma distribution is shown along with that for the three category point mass DFE in Figure 2.

**Table 4.** Bayesian estimates and 95% credible intervals for parameters of gamma distributions of negative and positive mutation effects (indexed by 0 and 1, respectively), under a two-sided gamma distribution models with the same or different means for negative- and positive-effect mutations. For example, *e̅*_1_ is the estimated mean of the gamma distribution of positive effect-mutations and *q*_1_ is their frequency.

| *Parameter* | *Model* | *Estimate* | *95% credible interval* |  |
| --- | --- | --- | --- | --- |
| $\beta$ | Two-sided gamma, same means | 0.32 | 0.26 | 0.70 |
| $\bar{e}$ | | 0.0049 | 0.0037 | 0.0070 |
| $q_1$ | | 0.48 | 0.39 | 0.58 |
| $\beta$ | Two-sided gamma, different means | 0.30 | 0.24 | 0.71 |
| $\bar{e}_0$ | | -0.0092 | -0.020 | -0.0060 |
| $\bar{e}_1$ | | 0.0021 | 0.0013 | 0.0032 |
| $q_1$ | | 0.84 | 0.73 | 0.90 |

**Figure 2.**
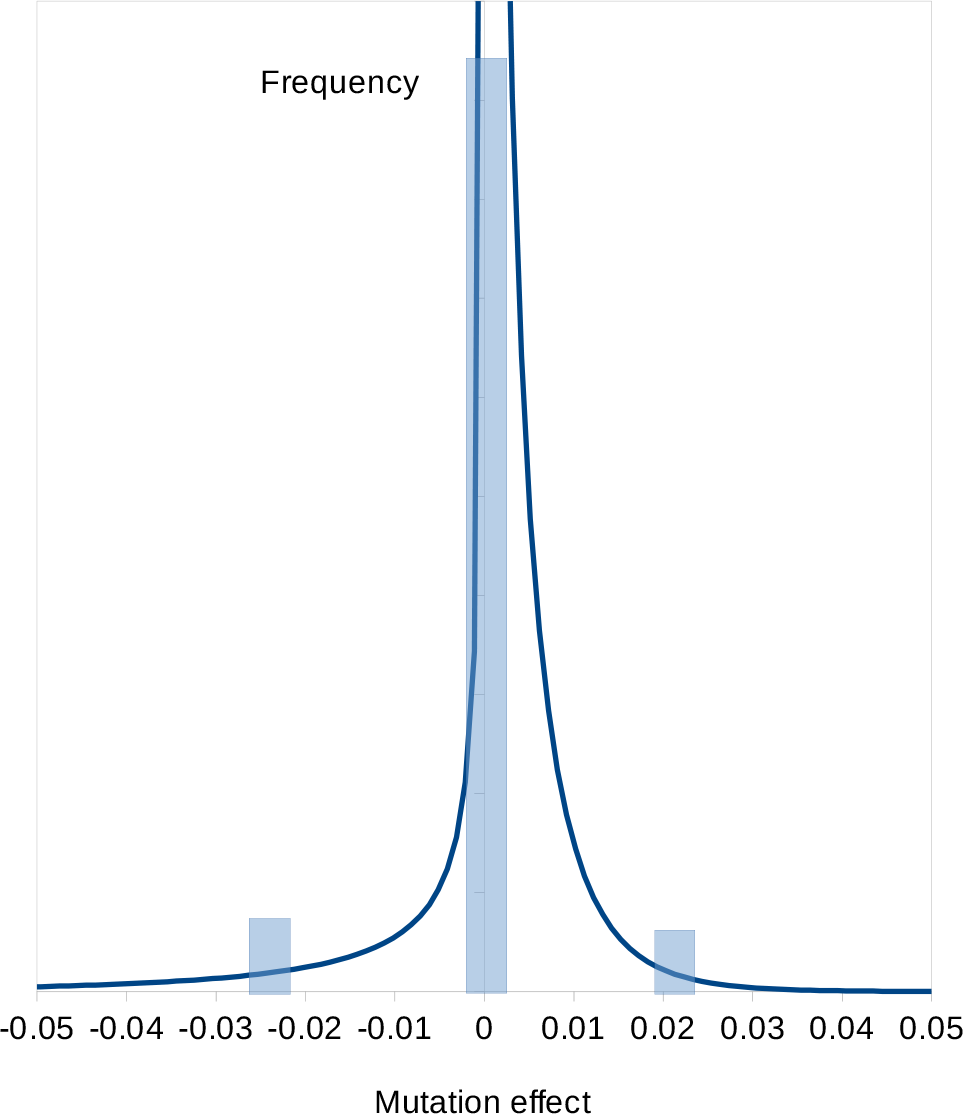
Inferred DFE assuming a two-sided gamma model (smooth line) and a point mass DFE for the three category model (transparent blue rectangles).

The credible intervals for the absolute means of negative- and positive-effect mutations do not overlap (Table 4), suggesting that the model with different means is more parsimonious that a model assuming a two-sided gamma distribution with the same means (Table 4; Figure S8). A model with different shape parameters for negative- and positive-effect mutations gives similar estimates for the mean effects and proportion of positive-effect mutations as the model with a single shape parameter, but stable estimates of the shape parameters could not be obtained, suggesting that this model is over-parameterised. We also analysed a more complex mixture model that, in addition to gamma-distributed negative- and positive-effect mutations, incorporated a class of zero-effect mutations. This allows a discontinuous DFE that has a “spike” at zero. However, the modal estimate for the frequency of the zero-effect class was zero and its upper credible value was 1%, indicating that the two-sided gamma distribution captures the leptokurtic shape of the DFE adequately.

### Relationships between estimated mutation effects and mutation types

To investigate whether mutations in certain mutation classes (such as exonic/non-exonic) are more or less likely to be associated with fitness, we calculated the effect of each mutation (as the posterior mean) under the two-sided gamma distribution model and then computed the difference between the average squared effects for mutations in mutually exclusive annotation classes. We examined average squared differences, because the additive variance contributed by a mutation is proportional to its squared effect. The results are negative in the sense that there are no statistically significant relationships for any of the mutation types tested (Table 5).

**Table 5.** Average squared effects of mutations (x1000) of certain mutation type classifications estimated under the two-sided gamma distribution model. For example, in the row labelled “SNP v Indel”, 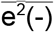 and 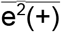 are the average squared effects for SNP and indel mutations, respectively. *P*-values for the difference between the squared effects of mutations were obtained by bootstrapping mutations 1,000 times.

| Mutation type | $\overline{e^2(-)}$ | $\overline{e^2(+)}$ | *P*-value |
| --- | --- | --- | --- |
| SNP v Indel | 0.074 | 0.073 | 0.84 |
| Non-exonic v Exonic | 0.061 | 0.083 | 0.15 |
| Non-intronic v Intronic | 0.081 | 0.059 | 0.17 |
| Non-intergenic v Intergenic | 0.074 | 0.067 | 0.92 |

## Discussion

In this paper, we integrate information on the fitness of MA lines, ancestral lines and crosses between MA lines and their ancestors with the complement of mutations carried by each line or cross. By crossing MA lines with their ancestors, each RL is expected to contain a different complement of mutations, which can be determined by genotyping. If there is sufficient replication, it is possible to estimate the individual phenotypic effects of mutations. The total number of mutations genotyped in the six MA lines studied was 386, however, implying that the effects of most mutations must be very small, and estimation of a fixed effect for each mutation is inappropriate. We therefore developed a random effects model, fitted a mixture of distributions using MCMC, and obtained Bayesian estimates of the parameters of the distributions. We investigated models in which each mutation is assigned to one of a number of classes of fitness effects (which includes a class with zero effect) or we assume that mutation effects are drawn from a mixture of gamma distributions. Our approach has similarities to the Bayesian mixture model method BayesR (Moser et al 2015) developed to estimate the distribution of SNP effects in genome-wide association studies. BayesR simultaneously analyses all informative SNPs (we likewise include all mutations) and fits a mixture of distributions of SNP effects, including a zero-effect class. Specifically, BayesR estimates the relative frequencies of the zero-effect class and a mixture of normal distributions of SNP effects with fixed variances. In this respect, BayesR differs from our method, where we estimate discrete categories of effects or gamma distribution parameters as variables in the model, and we also simultaneously estimate the frequencies of mutations in the different effects categories or gamma distributions.

Previous approaches to infer the distribution of fitness effects for spontaneous mutations (the DFE) using data from MA experiments have compared the distributions of estimated trait values for MA lines and unmutated controls. The simplest approach is the Bateman-Mukai method (Bateman 1959; Mukai 1964), which uses the changes of trait mean and genetic variance between MA lines and unmutated controls to estimate a genomic mutation rate parameter (*U*, the frequency of mutations with an effect on the trait) and the average effect of a mutation (*e̅*), while assuming that all mutations have the same effect. The information that can be obtained by the Bateman-Mukai method, and other approaches that use the full distribution of MA line phenotypic values (Keightley 1994; García-Dorado 1997; Shaw et al 2002), is extremely limited, however (reviewed by Halligan and Keightley 2009). The limitation arises because the genomic mutation rate and *e̅* are confounded with one another under the Bateman-Mukai approach, so the DFE and *U* are also confounded, and there is little information to distinguish between alternative models for the DFE if the effects of mutations are assumed to vary (Keightley 1998).

For five of the six CC-2931 MA line crosses, there is a negative relationship between growth rate and the number of mutations carried by a RL, although in some cases the relationship is very weak. This result is broadly consistent with the tendency for most *C. reinhardtii* MA lines to have lower growth rate than their ancestors (Morgan et al 2014; Kraemer et al 2016), and with Kraemer et al (2017), who generally observed negative relationships between fitness measured in competition with a marked strain and the numbers of mutations carried for MA lines of several genetic backgrounds. Kraemer et al (2017) also attempted to estimate a multi-category DFE based on the relationship between mutation number and fitness, but the amount of information available was limited, principally because there were only 10-14 MA lines tested of each genetic background. Here, we have characterized 1,526 RLs and a large number of combinations of genotypes, and therefore expect this design to be more powerful for inferring properties of the DFE than previous approaches that analysed individual MA lines.

We first investigated models in which mutation effects fall into discrete categories, including a zero-effect category. Under a two category model, there is a strong signal of growth rate-reducing mutations (estimated effect ~ −3%), consistent with the overall negative effect of spontaneous mutations we previously observed. The majority of mutations (~96%) are, however, allocated to the zero-effect class. Under a three class model, most mutations are also allocated to the zero-effect class, there is a negative-effect category with similar fitness effect and frequency as in the two class model, and a third category of positive-effect mutations (effect ~+2% on fitness). The frequency of positive-effect mutations is ~6%, but the credible interval is very wide. We then analysed a two-sided gamma distribution model, in which there are different means for the distributions of positive- and negative-effect mutations. Arguably, this is more realistic than the multi-category model, which assumes that mutation effects are invariant within a category. Consistent with the results from the analysis of the model with three discrete categories, there are both negative-and positive-effect mutations, and the proportion of positive-effect mutations is surprisingly high (~80%). The distributions for negative- and positive-effect mutations are highly leptokurtic (i‥e., the estimate of the shape parameter is ~0.3), and the absolute means of the distributions are both <1%, reflecting the concentration of density around zero. It appears that the effects of positive mutations are smaller than negative mutations, and the amount of mutational variance contributed by positive-effect mutations is ~20% that of negative-effect mutations. The fit of the estimated two-sided gamma distribution of effects is compared to the frequency distribution of the estimated effects of the individual mutations in Figure 3. Overall, the fit to the expected distribution is reasonable, although it appears that the inferred distribution may under-fit negative-effect mutations. There is one mutation with a positive effect of +5% (a G→C mutation in the 3'UTR of a gene on chromosome 6 of unknown function) and several mutations with absolute negative or positive effects >1%. The annotations associated with the 10 mutations with the highest absolute effects (i.e., the most extreme 2.5%) are shown in Table S6. There is no significant enrichment of any annotation we tested for these most extreme effects (or for the most extreme 5%; data not shown).

**Figure 3.**
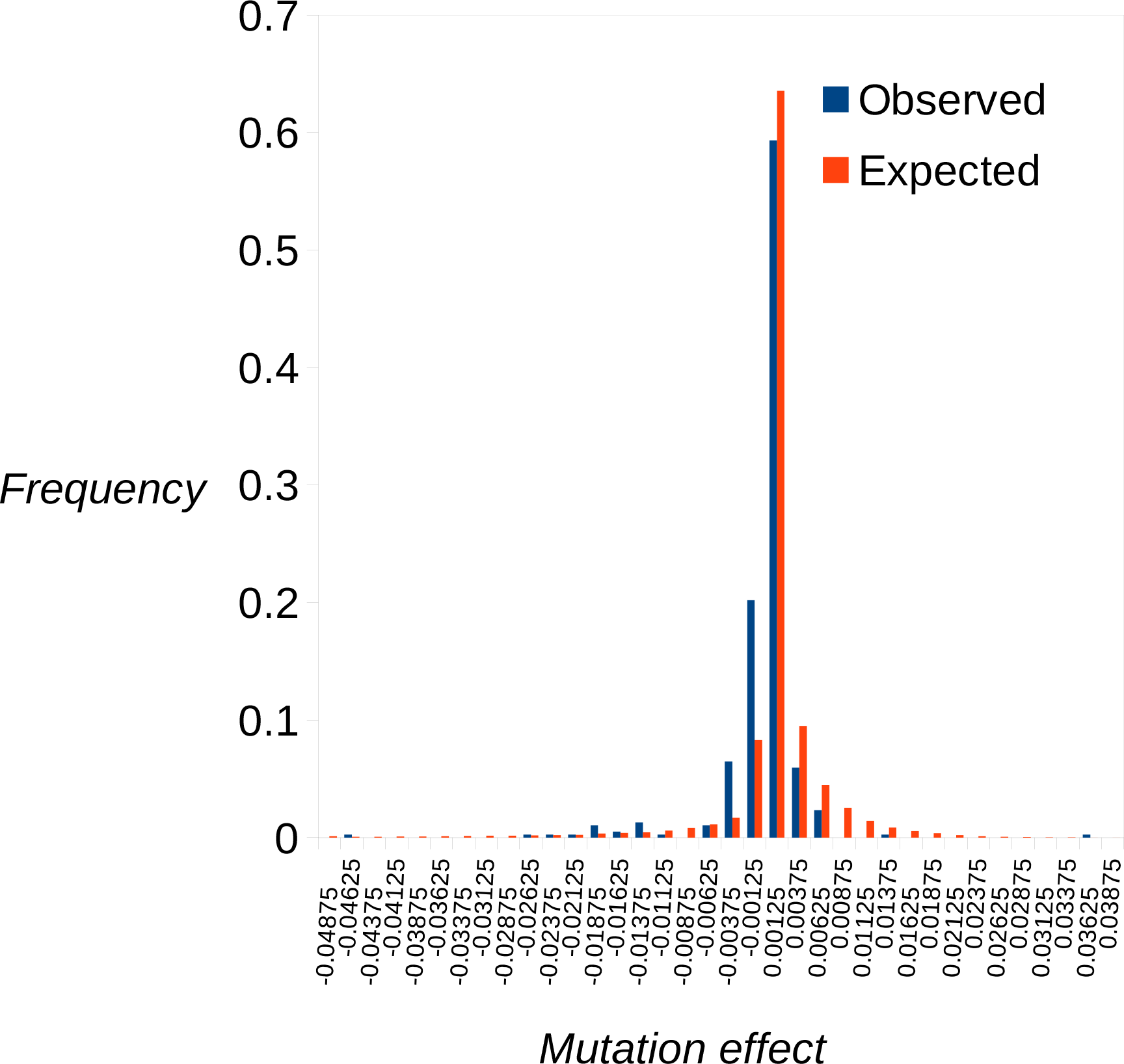
The estimated reflected gamma distribution of effects (expected) compared to the distribution of posterior mean estimates for the effects of the individual mutations (observed).

Why are we seeing a high proportion of positive-effect mutations? One possibility is that mutations that increase fitness are common in natural populations, and this is reflected in MA experiments (Shaw et al 2003; Rutter et al 2018). An alternative view is that deleterious mutations predominate in nature, principally because organisms are well adapted to the environments they typically experience (Keightley and Lynch 2003). Consistent with this, functional elements of the genome are typically conserved (Graur et al 2013), and analysis of the frequency of amino acid and synonymous polymorphisms within populations suggests that advantageous amino acid mutations are infrequent (Schneider et al 2011). A second possible explanation is that the algae were assayed in an environment which the species does not encounter in the wild, and some mutations that are deleterious in nature increase growth rate in the laboratory. A third possibility is that natural selection could not be prevented during the MA experiment, and there was either positive selection for mutations increasing growth or negative selection acting on mutations decreasing growth rate. This could take the form of between colony selection, if the fastest growing colonies were picked preferentially. Alternatively, there could be within colony selection, if new advantageous mutations occurring during colony expansion rise in frequency, or new deleterious mutations are removed during colony expansion. The effective population size was ~7 (Morgan et al. 2014), and we infer that few mutations have positive effects >10%, so any substantial selection for positive-effect mutations seems unlikely. On the other hand, deleterious mutations with effects >10% (including lethal or near-lethal mutations) would be under-represented.

The inferred DFE is is highly leptokurtic; many mutations have a very small effect and their is a long tail of large effect mutations. Under the reflected gamma distribution model the shape parameter of the distribution of negative- and positive-effect mutations is ~0.3. This is close to the value assumed by Kimura (1979, 1983) when analysing the nearly neutral model of molecular evolution, and is therefore consistent with the observation that amino acid variation is relatively insensitive to *N*_*e*_ (Ohta 1977; Kimura 1983), since a high proportion of sites in the Chlamydomonas genome are in protein-coding exons. It is also relevant to the narrow range of variation observed at synonymous and noncoding sites (Leffler et al 2012), if such sites become effectively selected in populations of large effective size. Such a leptokurtic distribution also has implications for the response to artificial selection and maintenance of variation for quantitative traits. If mutation effects are drawn from a leptokurtic distribution, then the response from new mutations builds up quicker than under the infinitesimal model, but is more variable, since response depends on the chance appearance and fixation of mutations with large effects (Hill 1982). A weak relationship between genetic variance for fitness or a correlated trait and *N*_*e*_ is also predicted (Keightley and Hill 1990).

To our knowledge, our approach of crossing MA lines to their ancestors and genotyping and phenotyping the crosses has not been previously attempted. It is related to that applied to induced mutations in RNA viruses (Sanjuan et al 2004) and in mismatch repair–deficient *E. coli* (Robert et al 2018). A limitations of our approach is that some mutations we previously identified by whole-genome sequencing (Ness et al 2015) were not amenable to genotyping. Specifically, some classes of mutations, such as large indel events or transposable element insertions, were not detectable by our short-read sequencing study, or may have occurred in regions that could not be aligned to the reference genome (Ness et al 2015). The approach is limited by the precision of phenotyping for mutations with small effects on growth rate. In general, laboratory-based measurements of mutation effects on fitness have been limited to those stronger than 10^−3^ (Gallet et al 2012). Mutations with effects of this magnitude or below will be allocated to the zero-effect category under the discrete class model or will have estimated effects close to zero under the two-sided gamma distribution model. Such mutations might be effectively selected in natural populations, however. The approach therefore has the capability of informing about mutations which may be under such strong selection in nature, and as such rarely segregate in natural populations. Other approaches that focus on the frequency distribution of segregating polymorphisms (Eyre-Walker et al 2006; Keightley and Eyre-Walker 2007; Boyko et al. 2008; Tataru et al 2017) inform about weakly selected mutations and therefore complement the present approach.

## Materials and Methods

### Mutation accumulation lines and the ancestral strain

Production and sequencing of MA lines of six strains of *C. reinhardtii* has been described previously (Morgan et al. 2014, Ness et al. 2015). Here, we focus on MA lines derived from CC-2931, a strain first sampled in Durham, North Carolina in 1991 that has a typical mutation rate among several strains we investigated (Ness et al. 2015) and decreasing mean fitness with increasing mutation number (Kraemer et al. 2017).

The MA lines and their ancestral strain are of the same mating type (mt-), so we first produced a “compatible ancestor” to which the MA lines could be crossed. This was done by backcrossing CC-2931 to a strain of the opposite mating type (CC-2344, mt+) for 13 generations with the aim of producing a strain identical to CC-2931, except for the region around the mating type locus on chromosome 6. Genome sequencing of the compatible ancestor (using the method of Ness et al. 2015) unexpectedly revealed, however, non-CC-2931 regions not only on chromosome 6 but also on chromosomes 4, 5, and 16 (Figure S9, S10) comprising a total of 7.6% of the genome and leaving 13 pure CC-2931 chromosomes. We dealt with this issue by including markers for these regions as factors in the analyses (see below).

### Generation of first generation recombinant lines (RLs)

For each MA line, we set up nine independent matings with the compatible ancestor, and collected 32 recombinant lines (RLs) from each to obtain a total of 288 RLs per MA line. Matings were set up by inoculating cultures for both parents into 200 µl of liquid Bold's medium (Bold 1942), and incubating these under standard growth conditions (23°C, 60% humidity, constant white light illumination) while shaking at 180 rpm for four days. Nitrogen-free conditions are required to trigger mating in *C. reinhardtii* (Sager & Granick 1954), so we centrifuged the cultures (3500 g, 5 minutes), removed the supernatant, and added 200 µl of nitrogen-free liquid Bold's medium. We then mixed 50 µl each of MA line and compatible ancestor cultures and incubated the matings for approximately 24 hours under standard growth conditions to allow zygotes to form at the surface. The zygote mats were transferred to Petri dishes containing Bold's agar and incubated in the dark for five days to allow zygote maturation. To kill any vegetative cells associated with the zygote mats, the Petri dishes were exposed to chloroform for 45-60 seconds. Subsequently, the Petri dishes were incubated under standard growth conditions until the matured zygotes had germinated. As controls for the chloroform treatment, 30 µl of both of the unmated parents of each of the mating reactions were subjected to the same procedure, and the respective mating reaction was discarded if any growth was observed. After successful germination, 2 ml of liquid Bold's medium was added to the Petri dishes to allow the germinated cells to go into suspension. The suspensions were then diluted and spread onto new Petri dishes containing Bold's agar and incubated under standard growth conditions until individual colonies had grown sufficiently to be picked. Initially 36 individual clones representing individual RLs were picked from each mating and transferred into 200 µl liquid Bold's medium and incubated under standard growth conditions while shaking at 180 rpm for three days. Finally, 32 of the 36 picked RLs were transferred onto Bold's agar in 7 ml bijou containers for long-term storage.

### Sample preparation for DNA extraction and genotyping

We used the competitive allele-specific PCR (KASP^TM^, Kompetitive Allele Specific PCR) technology to genotype the RLs of each MA line, the corresponding MA line, the compatible ancestor and the original unmutated ancestral strain (CC-2931) at the locations of the mutations previously reported for the MA lines (Ness et al. 2015). For allele-specific primer design, DNA regions of 1,000 base pairs (bp) surrounding each mutation were extracted from the *C. reinhardtii* reference genome (strain CC-503; version 5.3; Merchant et al. 2007). The regions were then corrected to match the consensus sequence of the CC-2931 MA lines.

For DNA extraction, we obtained cell pellets of at least 50 mg as follows. We inoculated the RLs and ancestors into 200 µl of liquid Bold's medium and incubated these under standard conditions with shaking at 180 rpm for four days. The cultures were then transferred to individual wells of 6-well plates filled with 6 ml of Bold's agar and incubated under standard conditions until a thick lawn had grown. Cells were then scraped off, transferred to 2 ml tubes and frozen at −70 °C. DNA samples were extracted from the frozen cell pellets and genotyped by LGC Genomics (http://www.lgcgenomics.com) using the sequences flanking each mutation of interest.

In addition to genotyping the known mutations, we genotyped markers that distinguish the mating types and the non-CC-2931 regions (Figure S9). For the mating type locus we designed markers matching loci specific to the two mating types, the *fus1* locus for the mt+ mating type and the *mid* locus for the mt-mating type. For the non-CC-2931 regions we included markers for sites that differed between the two strains within these regions.

### Determination of RL mating types by crossing

In addition to using genetic markers, we determined mating type using crosses. In separate mating reactions, we mated each RL with the ancestral strain and with the compatible ancestor, using a modification of the mating protocol described above, in which we extended the incubation period for the mating reaction under standard growth conditions to approximately 48 hours and then incubated plates in the dark for 5 days. To kill vegetative cells, we then incubated the plates for 5 hours at −20 °C, added 100 µl of a Bold's medium containing twice the amount of nitrogen as Bold's medium, and incubated the plates under standard growth conditions while shaking at 180 rpm until zygotes had germinated. We assigned mating type for each RL based on the combined results of the mating test and the mating type genotyping test. If one test failed, we used the result of the other. If the tests disagreed or both failed, a mating type was deemed not assignable and was recorded as missing data.

### Measurement of growth rate

To generate growth curves for the individual RLs and their parents (i.e., the corresponding CC-2931 MA line and the compatible ancestor), we inoculated each of these separately into individual wells of 96 well plates containing 200 µl of liquid Bold's medium. Each plate contained samples from 58 RLs, all derived from the same MA line and their parental lines. We allocated lines randomly among the 60 central wells to avoid plate edge effects (Morgan et al. 2014) and filled the outer wells with 200 µl of medium to maintain humidity and reduce evaporation in the central wells. All plates were initially incubated for four days under standard conditions. On day 4, we transferred 2 µl of each culture to the corresponding well on a new 96-well plate filled with 198 µl liquid Bold's medium to start the growth assay. As an estimate of cell density, we measured absorbance at 650 nm every 12 hours over a period of 96 hours. We repeated this complete procedure twice in order to have two temporally independent replicates for each RL.

Maximum growth rate can be estimated from each growth curve as the slope of the linear regression of the natural log (ln) of absorbance on time during the exponential phase of growth. Unfortunately, the start and duration of exponential growth varied between growth curves, so we were unable to simply estimate growth over the same time window for each growth curve. Instead we used the following procedure. For each growth curve we generated a number of 48 hour time windows which spanned 4 measurements in our growth curves. The first started at 12 hours and ran to 60 hours, the second started at 24 hours and ran to 60 hours and so on until we had all possible windows up until 96 hours. For each window we then carried out a regression of ln absorbance on time. The slope of this regression line gives us an estimate of the rate of increase during this time period, whilst the proportion of the total variation in growth rate explained by the linear regression on time (the *R*^2^ value) gives an estimate of how well the linear relationship fits the data. We carried out this procedure for windows of 60 hours (5 time points), 72 hours (6 time points) and 84 hours (7 time points). We then excluded any windows for which the fit of the linear model was not adequate (*R*^2^ < 0.75). We then examined the slope estimates from each of the remaining windows and used the highest estimate as our measure of maximum growth rate for that growth curve. Visual inspection of the fitted lines on the time series showed that this procedure was effective in identifying the period of maximum growth for the variety of observed growth trajectories. For a total of 8 RL replicate time series measures, an adequate fit was not achieved for any of the time windows due to extremely unusual growth trajectories, and these were excluded from further analysis (Table S1).

### Data processing and preparation

Mutations that were invariant across all samples were considered as artifactual and excluded. We also excluded mutations that were in complete linkage with either the mating type locus or one or more marker from the non-CC-2931 regions (Table S2). In the case of 21 mutations, only one of the two allele-specific primers worked successfully, and consequently no genotype information on the mutation was available for about 50% of the RLs. We corrected such mutations by changing the missing genotype to the non-amplified allele (Table S7). We excluded RLs for which genotypes of more than 10% of mutations were missing and/or for which more than 5% of mutations were assigned as heterozygous. *C. reinhardtii* is haploid, and multiple heterozygous calls suggest that the RL contains several different genotypes, and potentially cross-contamination. The rationale for setting these thresholds for missing data and heterozygous calls came from plotting the distributions of percentages of missing data and heterozygous calls for the complete data set. Only a small number of RLs have more than 10% of missing mutations and/or have more than 5% of the mutations assigned as heterozygous (Figure S11).

After carrying out the above filtering steps, several missing genotypes remained, so we imputed as many as possible to facilitate analyses. We first assigned missing genotypes for cases where RLs of apparently identical haplotype originated from the same mating reaction. In a second step, we computed the squared measure of linkage disequilibrium between pairs of mutations (*r*²) (Charlesworth and Charlesworth 2010). We then examined the remaining mutations that have missing genotypes in turn. If *r*² between a mutation and its neighbouring mutation was above 0.7, we imputed its allelic state using the state of the neighbouring mutation. 1,982 (0.97%) of the total 205,351 data points across all MA lines were initially missing (=number of mutations x number of samples including all replicates of RLs and ancestors). With our imputation approach, we were able to impute 1,766 (89%) of them so that only 216 (0.11%) missing data points remained.

### Relationship between number of mutations and growth rate

To examine the relationship between RL growth rate and the number of mutations carried, we fitted a linear mixed model to the combined data set from all 6 MA lines and to the individual MA lines, with growth rate as the response variable and the number of mutations carried as a continuous linear predictor. To control for other sources of variation we also fitted mating type and all markers for the non-CC-2931 regions as fixed factors, and MA line, haplotype and growth assay plate as random factors. The significance of the number of mutations was examined by comparing models with and without this term, using a likelihood ratio test. The analysis was also done for specific mutation types (SNP, indel, exonic, intronic, intergenic). Models were fitted using the *lme4* (Bates et al. 2015) package in R (R Core Team 2018). The data along with the R code are provided in the supplementary online materials.

### Inference of the distribution of effects of mutations for growth rate

We developed a MCMC approach to infer the distribution of effects of mutations for growth rate, assuming two kinds of models. We modelled a discrete distribution in which each mutation’s effect fell into one of a number (*n_c_*) categories, and a two-sided gamma distribution allowing different parameters for the distributions of negative- and positive-effect mutations. To control for the effects of mating type and presence/absence of non-CC-2931 chromosomal regions, we estimated *n*_*f*_ two-level fixed effects. Normally distributed plate effects and residual effects and the overall mean were also fitted. When analysing a merged data set of all six MA line crosses, a different mean was fitted for each MA line and any RL with one or more missing genotypes was excluded.

The data for the RLs bred from one MA line are represented in Table 6. Let *n*_*b*_ be the number of RL observations and let *n*_*m*_ be the total number of mutations in the MA line. Mutations are encoded in a *n*_*b*_ x *n*_*m*_ matrix, **M**, whose elements (0 or 1) indicate the presence or absence of a mutation in a RL. The fixed effects associated with each observation are encoded in a matrix, **F**, of dimension *n*_*b*_ x *n_f_* with elements 0 or 1. Plate numbers (*n*_*p*_ levels) and phenotypic values associated with each observation are vectors **r** and **y**, respectively, both of dimension *n_b_*.

**Table 6.** Representation of the data from a MA line crossing experiment

| *Description* | *Symbol* | *Dimension* |
| --- | --- | --- |
| Number of observations | $n_b$ | Scalar |
| Overall number of mutations | $n_m$ | Scalar |
| Mutation matrix | $\mathbf{M}$ | $n_b \times n_m$ |
| Number of fixed effects | $n_f$ | Scalar |
| Fixed effects matrix | $\mathbf{F}$ | $n_b \times n_f$ |
| Number of plates | $n_p$ | Scalar |
| Plate number vector | $\mathbf{r}$ | $n_b$ |
| Phenotypic value vector | $\mathbf{y}$ | $n_b$ |

Each MCMC iteration, the state of one of the model’s variables (which are elements of vectors or scalars defined in Table 7-9) is proposed, then accepted or rejected based on change in log likelihood. Posterior distributions of the model variables provide Bayesian parameter estimates.

### Multi-category model

Under this model, one category of mutations has no effect on fitness, and the remaining categories have non-zero effects. The mutation category vector (**m**) specifies the category in which each mutation currently resides, the value zero signifying that a mutation is in the zero effect category (Table 7). Elements of **m** are proposed by randomly picking an integer in the range 0 ‥ 1 – *n_c_*, which is different from the current value. State variables for the effects and frequencies associated with each category are encoded in vectors **e** and **q**, respectively. The first element (*e*_0_) of **e** is fixed at zero, since it is the effect of the invariant zero-effect class, and the first element of **q** is set to 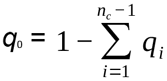, the frequency of the zero-effect class. Proposals for the remaining elements of **q** are random uniform deviates added to the current value. Changes to the values of all other variables are drawn from normal distributions with mean zero. The variances of the uniform and normal distributions of proposal deviates are adjusted during a burn-in phase so that about 25% of proposals are accepted.

**Table 7.** Variables in the MCMC model specific to the multi-category model.

| *Description of variable* | *Symbol* | *Dimension* | *Constraints on value* |
| --- | --- | --- | --- |
| Vector of mutation categories | **m** | 1..$n_m$ | Integer, 0.. 1 – $n_c$ |
| Vector of mutation effects | **e** | 0.. $n_c - 1$ | – |
| Vector of mutation frequencies | **q** | 0.. $n_c - 1$ | $\sum_1^{n_c-1} q < 1$ |

**Table 8.** Variables of the model common to the multicategory and two-sided gamma distribution models.

| *Description of variable* | *Symbol* | *Dimension* | *Constraints on value* |
| --- | --- | --- | --- |
| Vector of fixed effects | **f** | 1.. $n_f$ | – |
| Vector of plate effects | **p** | 1.. $n_p$ | – |
| Random plate effect variance | $V_p$ | Scalar | – |
| Overall mean | $\bar{y}$ | Scalar | – |
| Residual variance | $V_e$ | Scalar | – |

Proposals are accepted or rejected by applying the Metropolis-Hastings algorithm based on the log likelihood of the data and the priors (which are designed to be uninformative), given the parameter values. The overall log likelihood contains contributions from the numbers of mutations in different categories, their frequencies, the random plate effect, and each observation, which are considered independent. Let **v** be a vector of dimension 0 to 1 – *n*_*c*_ containing the numbers of mutations in each of the *n*_*c*_ categories in the current proposal, and *multinomial*(*n_c_*, **q**, **v**) be the probability of sampling **v** from a multinomial distribution parameter **q**. Let *normal*(*y*, *y̅*, *V_e_*) be the normal distribution probability density function for point *y* with mean *y̅* and variance *V_e_*. Let *g_i_* be the genotypic value of RL *i*, which is the sum of the effects of the mutations it carries. This is calculated from the set of mutations carried by the RL (specified in **M**), the categories into which these mutations fall (specified in **m**) and the effect associated with each category (specified in **e**):

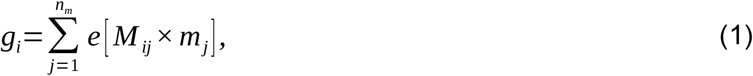

where the square brackets denote vector or matrix indexing, i.e., *e*[*x*] = *e*_x_. The overall log likelihood of the data is then:

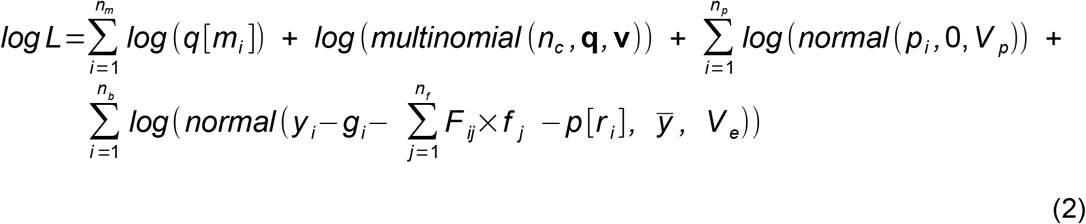

Note that the model with three categories (including a zero-effect category) is equivalent to a model with a mixture of two gamma distributions both with shape parameters → ∞ plus a zero-effect category (see below).

### Two-sided gamma distribution model

Under the two-sided gamma distribution model (whose variables are defined in Table 9), the current state of a mutation in the MCMC is defined by two variables. The first is whether the mutation has a negative or positive effect, encoded as 0 or 1 in vector **μ**. The second variable is the genotypic effect of the mutation, encoded in a matrix **E** of dimension [0‥1]x[1‥*n_m_*]. The current value of the element of **μ** selects the mutation’s current genotypic effect, i.e., for mutation *i* the genotypic value is *E*[*μ_i_*][*i*]. The frequencies of negative- and positive-effect mutation are encoded in vector **q**. The scale and shape of the gamma distributions for negative and positive-effect mutations are encoded in vectors **α** and **β**, respectively. Proposals for *q*_0_ are random uniform deviates added to the current value, such that 0 < *q*_0_ < 1 and *q*_1_ = 1 - *q*_0_. A proposal for an element of **μ** is 0 if the current value is 1 and *vice versa*. Changes to the values of all other variables are drawn from normal distributions with mean zero with adjustment during the burn-in as described above.

**Table 9.** Variables in the model specific to the two-sided gamma distribution model.

| *Description of variable* | *Symbol* | *Dimension* | *Constraints on value* |
| --- | --- | --- | --- |
| Vector of mutation sign indicators | $\boldsymbol{\mu}$ | $1..n_m$ | Integer, 0..1 |
| Matrix of mutation effects | $\mathbf{E}$ | $[0..1] \times [1..n_m]$ | Positive real number |
| Vector of frequencies of negative and positive-effect mutations | $\mathbf{q}$ | 0..1 | $0 < q_0 < 1$ |
| Vector of shape parameters | $\boldsymbol{\alpha}$ | 0..1 | Positive real number |
| Vector of scale parameters | $\beta$ | 0..1 | Positive real number |

Let gamma(*x*, *α*, *β*) be the gamma distribution PDF for point *x*, given scale and shape parameters *α* and *β*, respectively. Let **v** be a vector (with two elements indexed by 0 and 1) containing the numbers of mutations with negative and positive effects in the current proposal, and *binomial*(*q*_0_, *v*_0_) be the probability of sampling *v*_0_ negative-effect mutations, given that the frequency of negative-effect mutations is *q*_0_. Let *g_i_* be the genotypic value of RL *i* (the sum of the effects of the mutations it carries, as above). This is calculated from the set of mutations carried by the line (specified in **M**), the types into which these mutations fall (i.e., negative or positive specified in **μ**) and the effect of each mutation (specified in **E**):

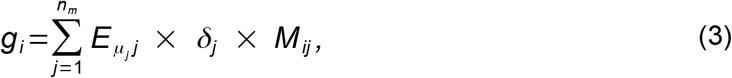

where *δ*_*j*_ takes the value −1 if *μ*_*j*_ is 0 and +1 if *μ*_*j*_ is 1. The overall log likelihood of the data is:

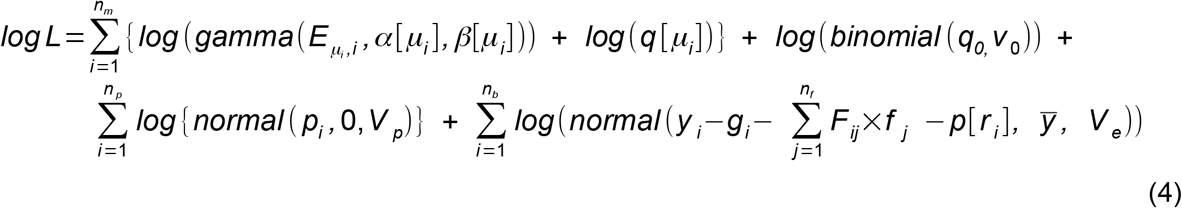

We considered models where the shape parameter of the gamma distributions for negative- and positive-effect mutations were the same or allowed to be different. We also implemented a somewhat more complex mixture model in which there are three categories of mutations: a zero-effect category and gamma-distributed positive and negative-effect mutations, as described above.

### Running the MCMC

MCMC runs started with a burn-in of 10^8^ iterations for multi-category models or 10^9^ iterations for two-sided gamma distribution models. Parameter values were then sampled every 10,000 iterations for 9×10^8^ iterations for multi-category models or for 5×10^9^ iterations for two-sided gamma distribution models. From each sampled iteration the vector of state variables was stored for generation of plots of parameter values against iteration number or posterior density plots. The posterior mode was taken as the parameter estimate, and 95% credible intervals computed on the basis of ranked parameter values. Priors for fixed effects, plate effects and the overall mean and variance were uninformative. The prior for mutation frequencies was a uniform distribution bounded by 0 and 1, and were therefore informative. Priors for the mutation effect parameters (under the multiple category model) were uniform in the range +/− 0.5 phenotypic standard deviations. Priors for the mean of the gamma distributions (under the two-sided gamma distribution model) were uniform in the range zero to 0.5 phenotypic standard deviations. Priors for the shape parameters of the gamma distributions were uniform in the range 0.1 to 100.

To check whether signals detected in the data were genuine, phenotypic values for fitness were permuted among backcross lines within plates without replacement. The distribution of estimates for parameters of interest obtained from such permuted data sets were computed. Significant estimates from the original data were expected to lie outside these distributions.

### Simulations

To check the method, simulated data sets with either two, three or four categories of mutational effects or a two-sided gamma DFE and 40 mutations in each data set were analysed as described above, while assuming the same model as simulated. In all cases posterior modes are close to the parameters values of the simulations (Figure S12-S15).

## Acknowledgements.

We thank Jarrod Hadfield for advice on the Bayesian mixture model and its implementation and Adam Eyre-Walker and Tom Booker for helpful comments on the manuscript. The work described was funded by a grant from the UK Biotechnology and Biological Sciences Research Council (BBSRC). This project has received funding from the European Research Council under the European Union’s Horizon 2020 research and innovation programme (grant agreement no. 694212).

## Author contributions.

Generated RLs [SAK, KBB, DM]. Carried out RL growth assays [SAK, KBB, DM, TSS, NC, PDK]. Carried out compatible ancestor generation, sequencing and analysis [JL, RWN, KBB]. Carried out RL genotyping and associated analysis and analysis of growth curves [KBB, TSS]. Carried out mating type assays [TSS]. Implemented and carried out MCMC analysis [PDK]. Wrote paper [PDK, KBB, NC]. Conceived study [RWN, NC, PDK].

## Supplementary Materials

### Supplementary figures

**Figure S1.**
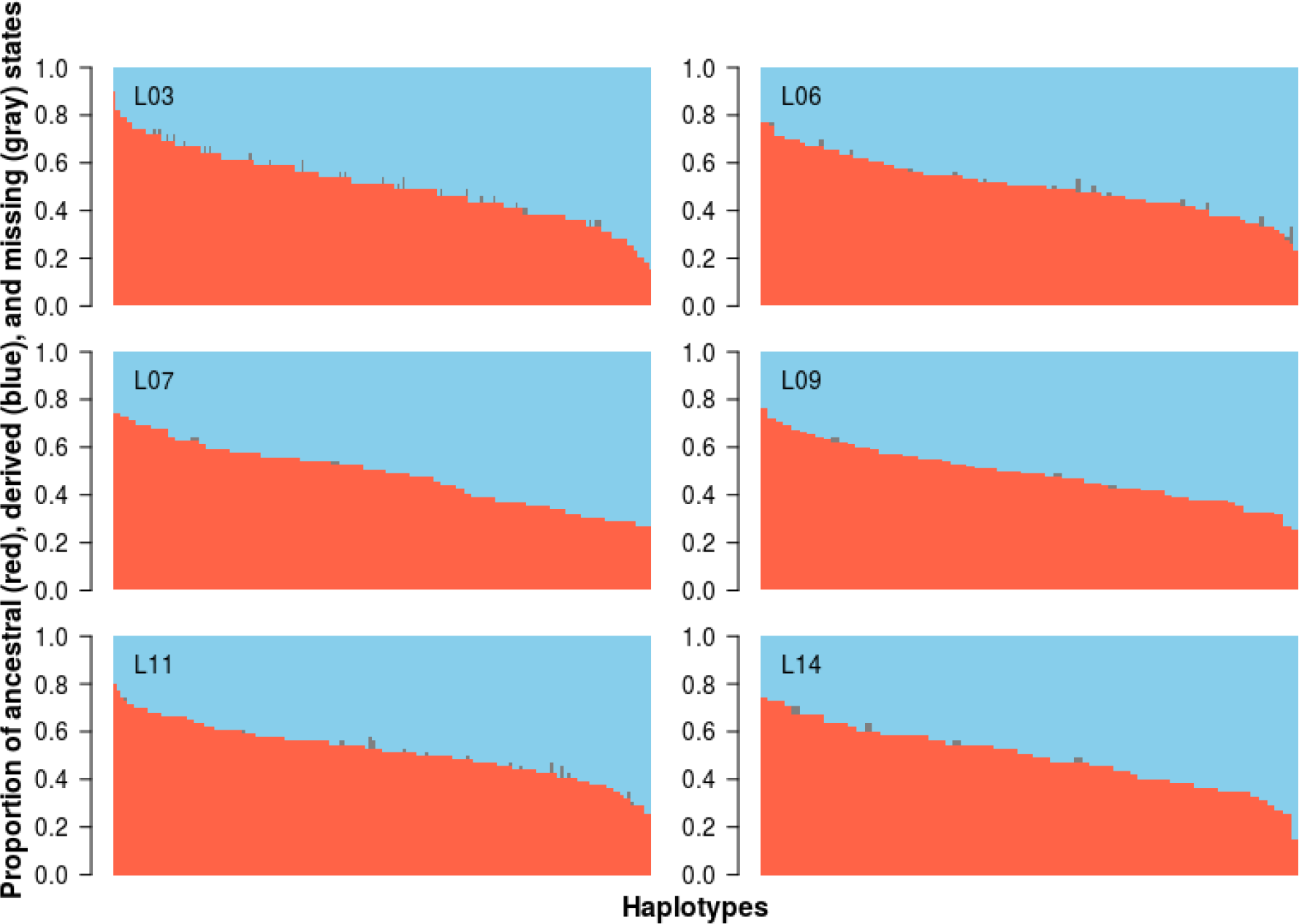
Proportion of ancestral (red), derived (blue) missing (grey) states at each mutated position for each haplotype. Haplotypes are sorted from left to right according to the proportion of ancestral states at the mutated positions.

**Figure S2.**
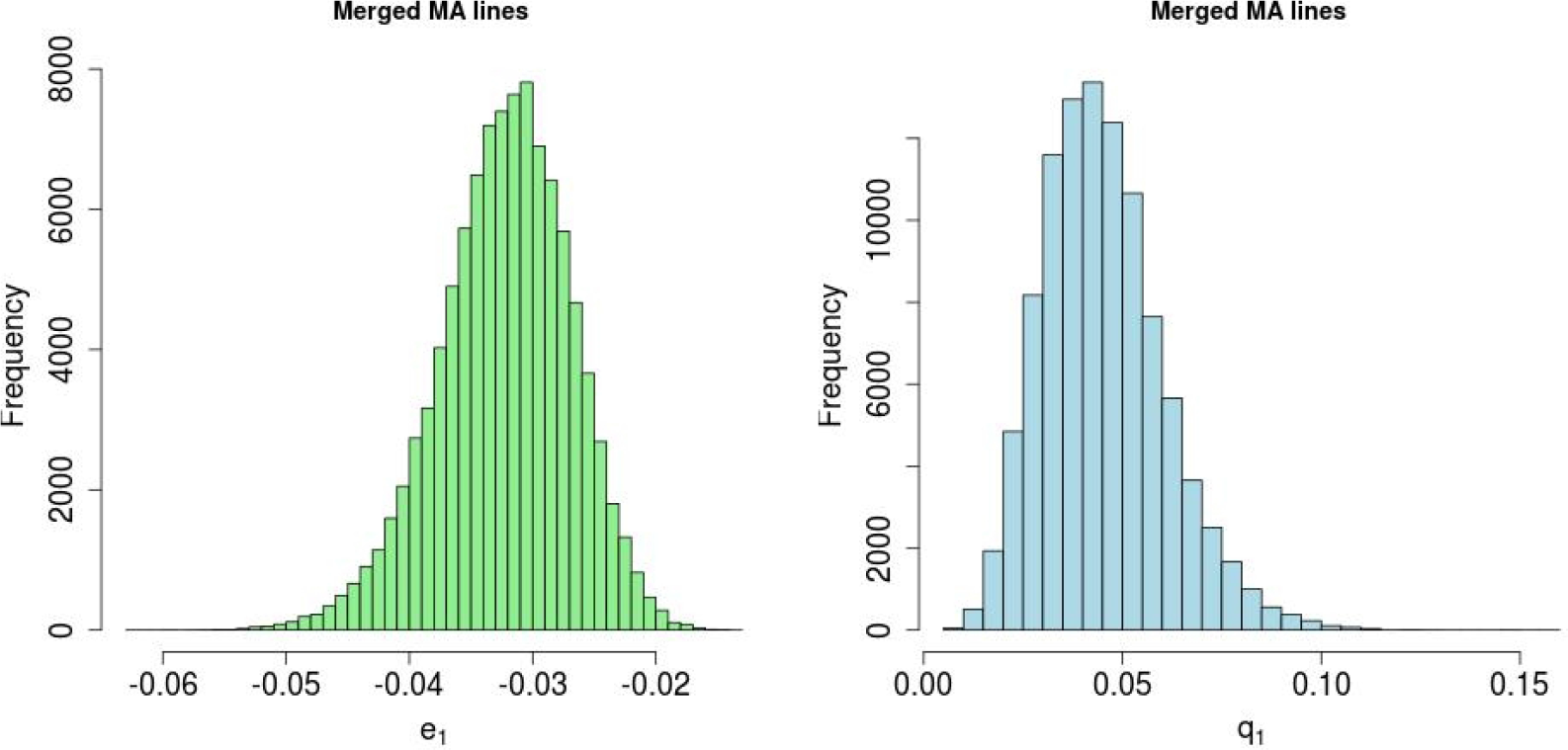
Results of MCMC analysis of combined data set of RLs from six MA lines, assuming a model with two categories of mutational effects one of which has an effect of zero. Bayesian posterior density plots are for parameters *e*_1_ and *q*_1_ (the effect and proportion of mutations in category 1, respectively).

**Figure S3.**
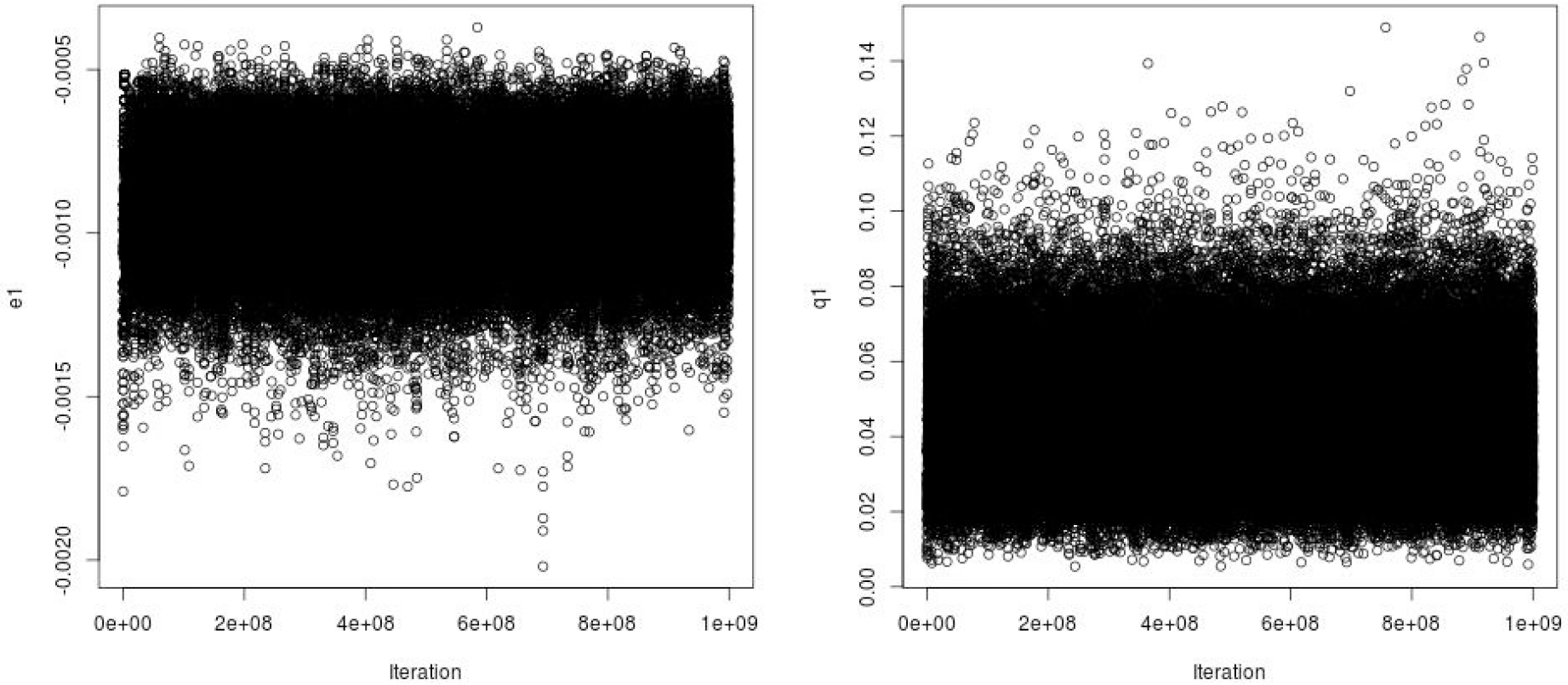
Values of sampled parameters *e*_1_ and *q*_1_ (effect and frequency of mutations) in MCMC run. The mutation effect is shown unscaled by the trait mean.

**Figure S4.**
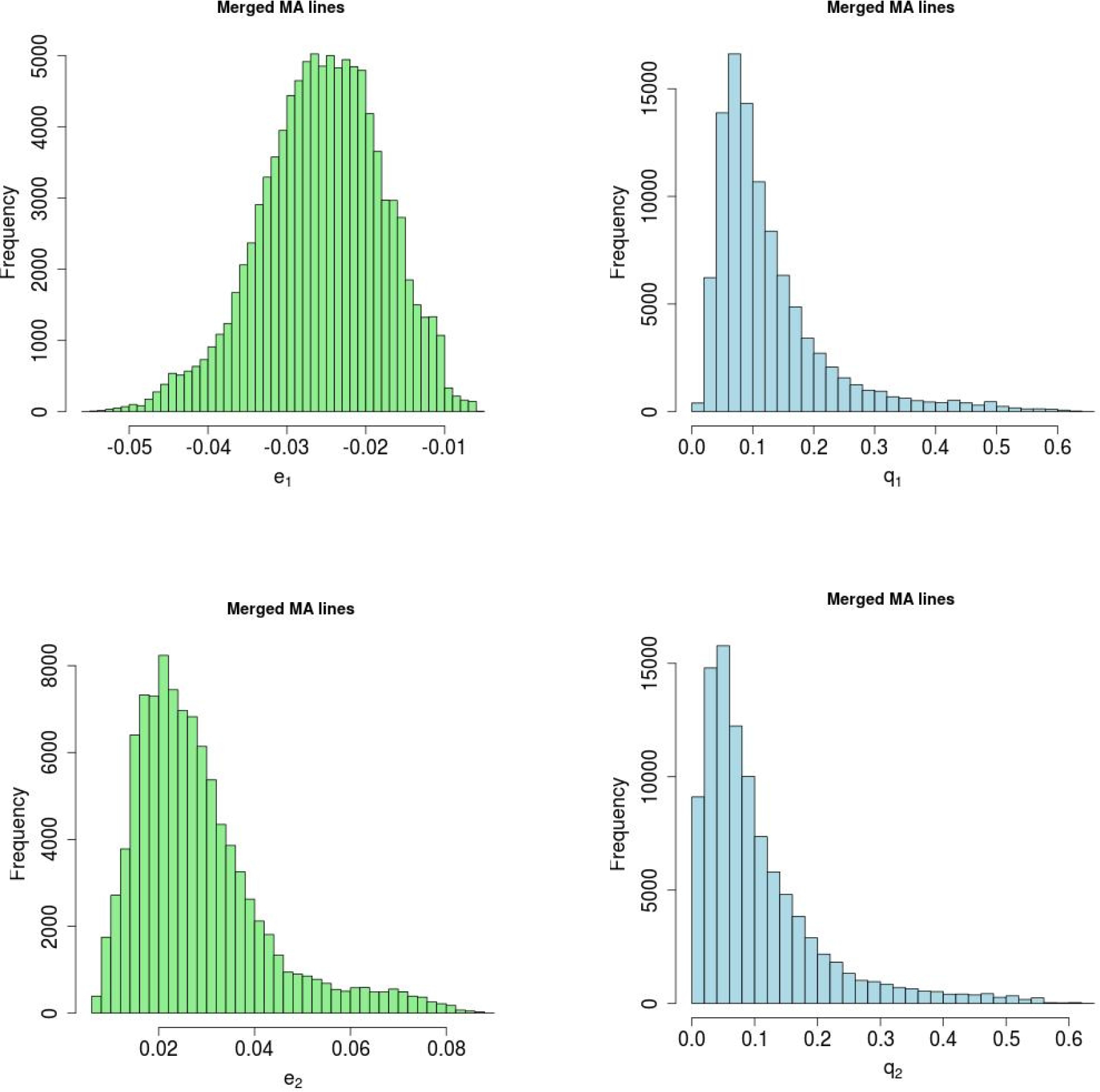
Results of MCMC analysis of combined data set of 6 MA line backcrosses assuming a model with three categories of mutational effects (including one category with an effect of zero). Bayesian posterior density plots are shown for *e* and *q* parameters (the effect of and proportion of mutations, respectively, in the two finite-effect categories).

**Figure S5.**
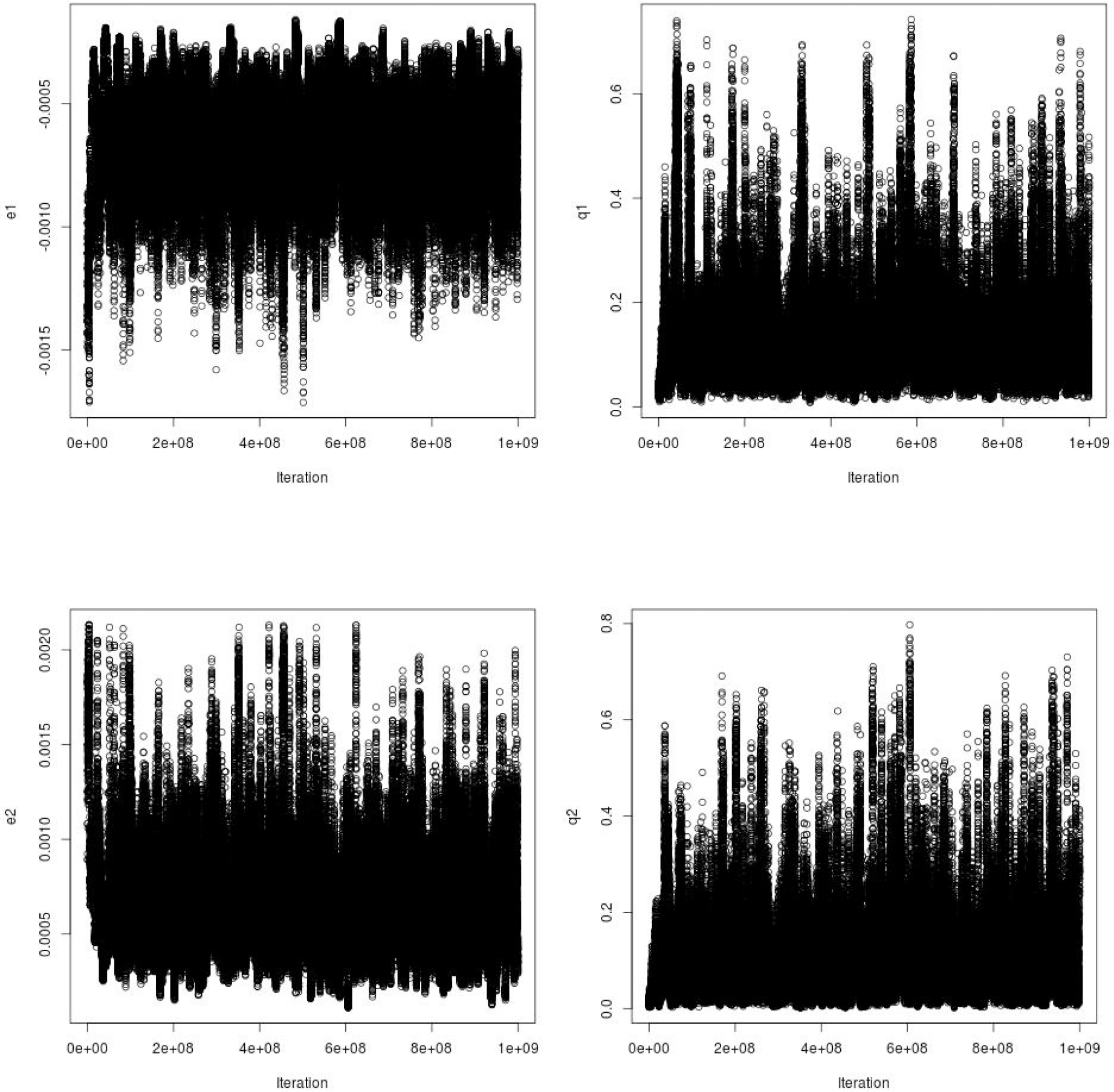
Values of sampled parameters *e* and *q* [effect and frequency for negative-(index 1) and positive-effect (index 2) mutations] in MCMC run. The mutation effects are shown unscaled by the trait mean.

**Figure S6.**
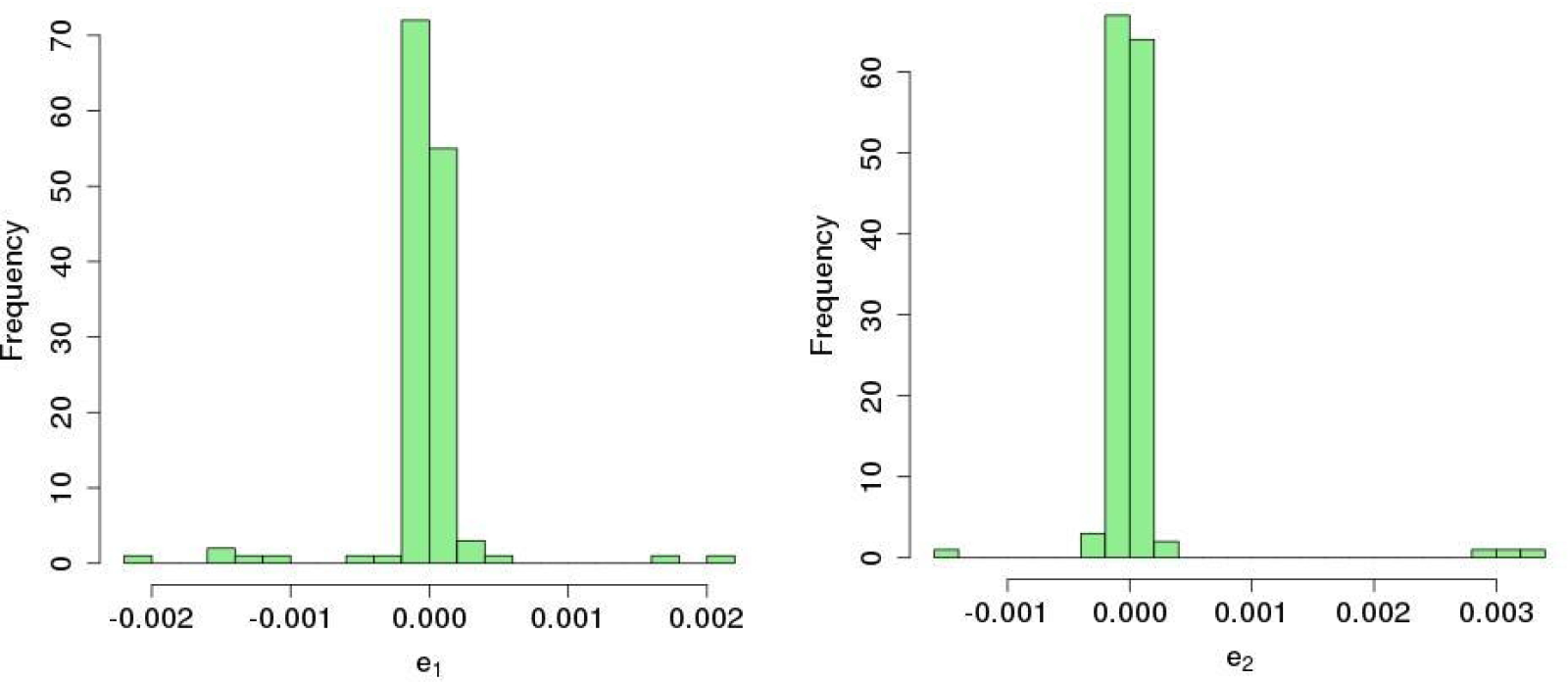
Distribution of posterior modal estimates for mutation effect parameters *e*_1_ and *e*_2_ obtained from analysis of datasets in which phenotypic values are permuted within plates with replacement under the three mutation category model.

**Figure S7.**
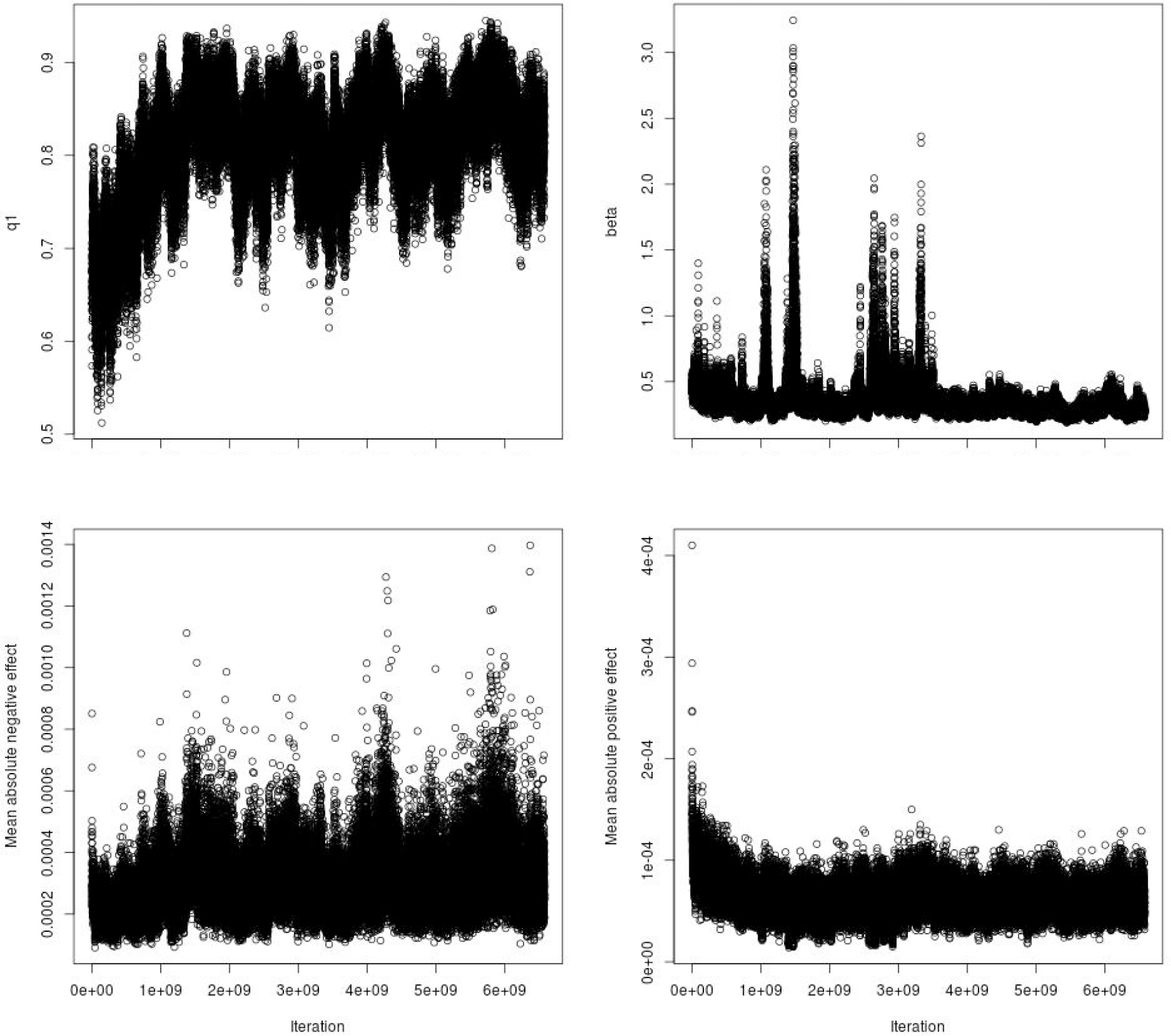
Values of sampled parameters *q*_1_ (frequency of positive-effect mutations), β (gamma distribution shape parameter) and means for negative and positive effect mutations in MCMC run for the two-sided gamma distribution with different means for negative- and positive-effect mutations. The mean mutation effects are shown unscaled by the trait mean.

**Figure S8.**
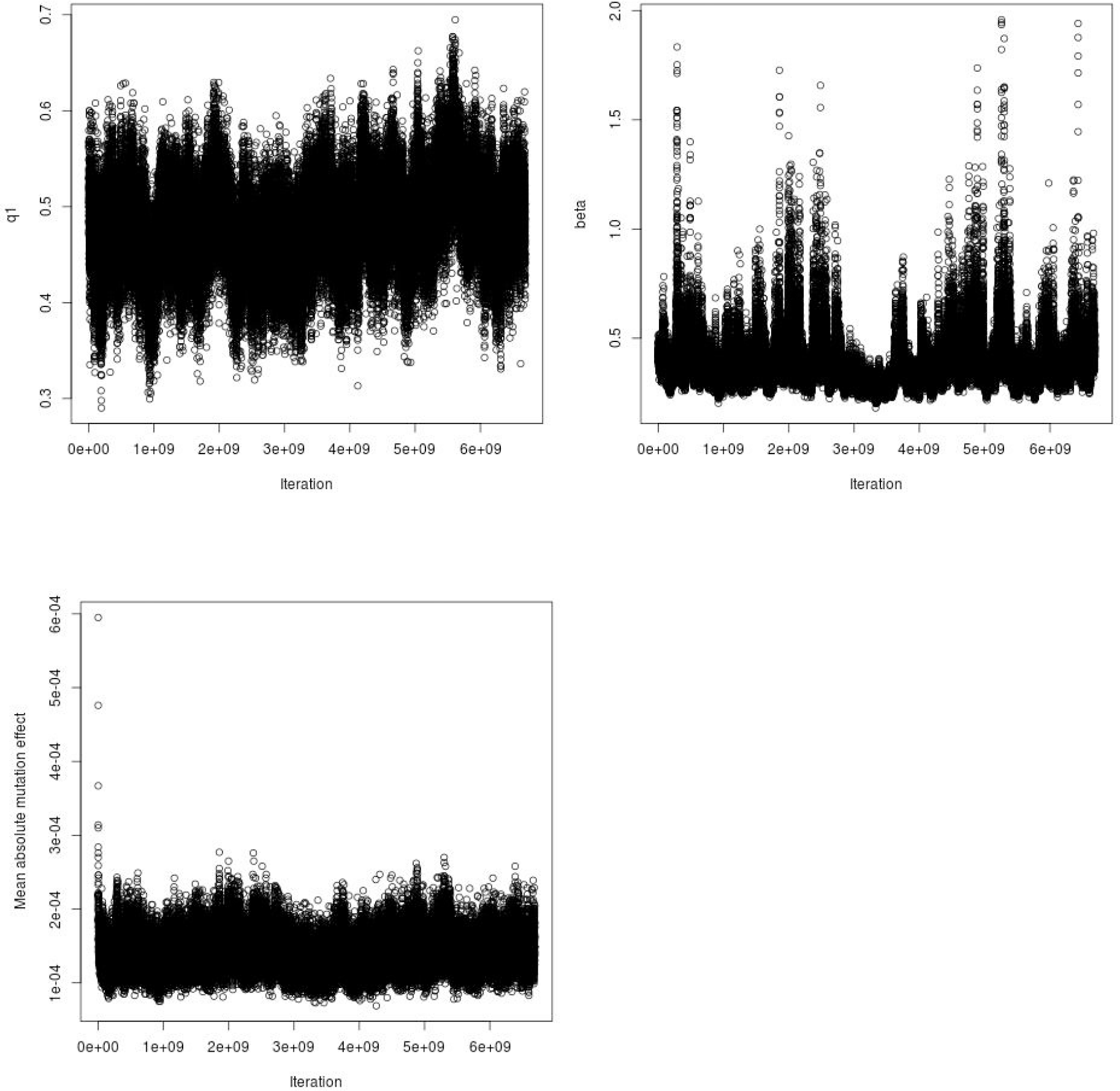
Values of sampled parameters *q*_1_ (frequency of positive-effect mutations), β (gamma distribution shape parameter) and mean absolute effect of mutations in MCMC run for the two-sided gamma distribution with the same means for negative- and positive-effect mutations. The mean absolute mutation effect is shown unscaled by the trait mean.

**Figure S9.**
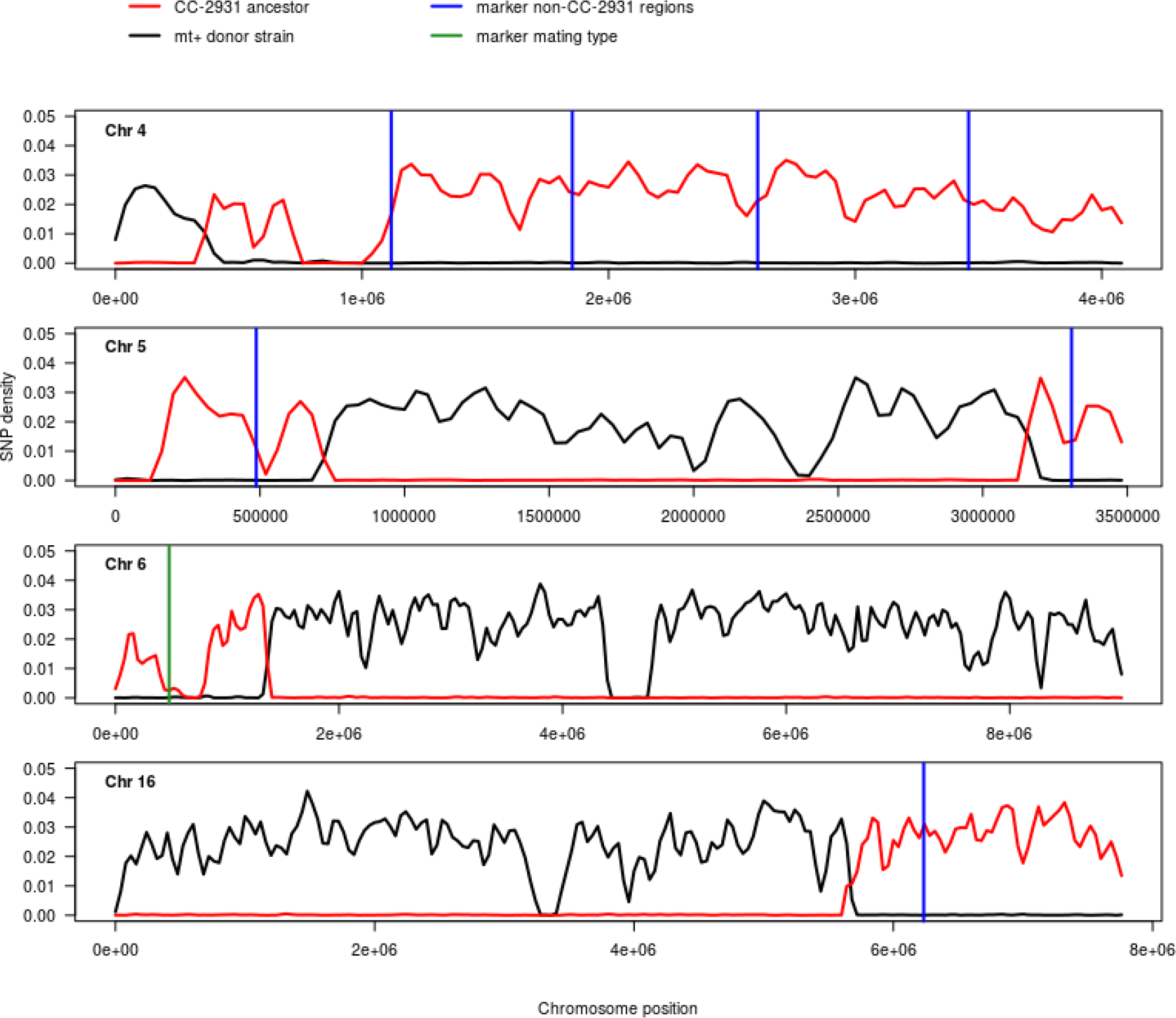
SNP densities along chromosomes 4, 5, 6, and 16 between the compatible ancestor for CC-2931 and its two ancestral strains. SNP densities were calculated for 80 kb windows along the chromosomes between the compatible ancestor and CC-2931 (red) and between the compatible ancestor and the mating type + donor strain (black). A mutation density of 0 indicates no genetic differences between the compatible ancestor and the strain it was compared to. The positions for the markers for the non-CC-2931 regions (blue) and for the mating type marker (green) are indicated.

**Figure S10.**
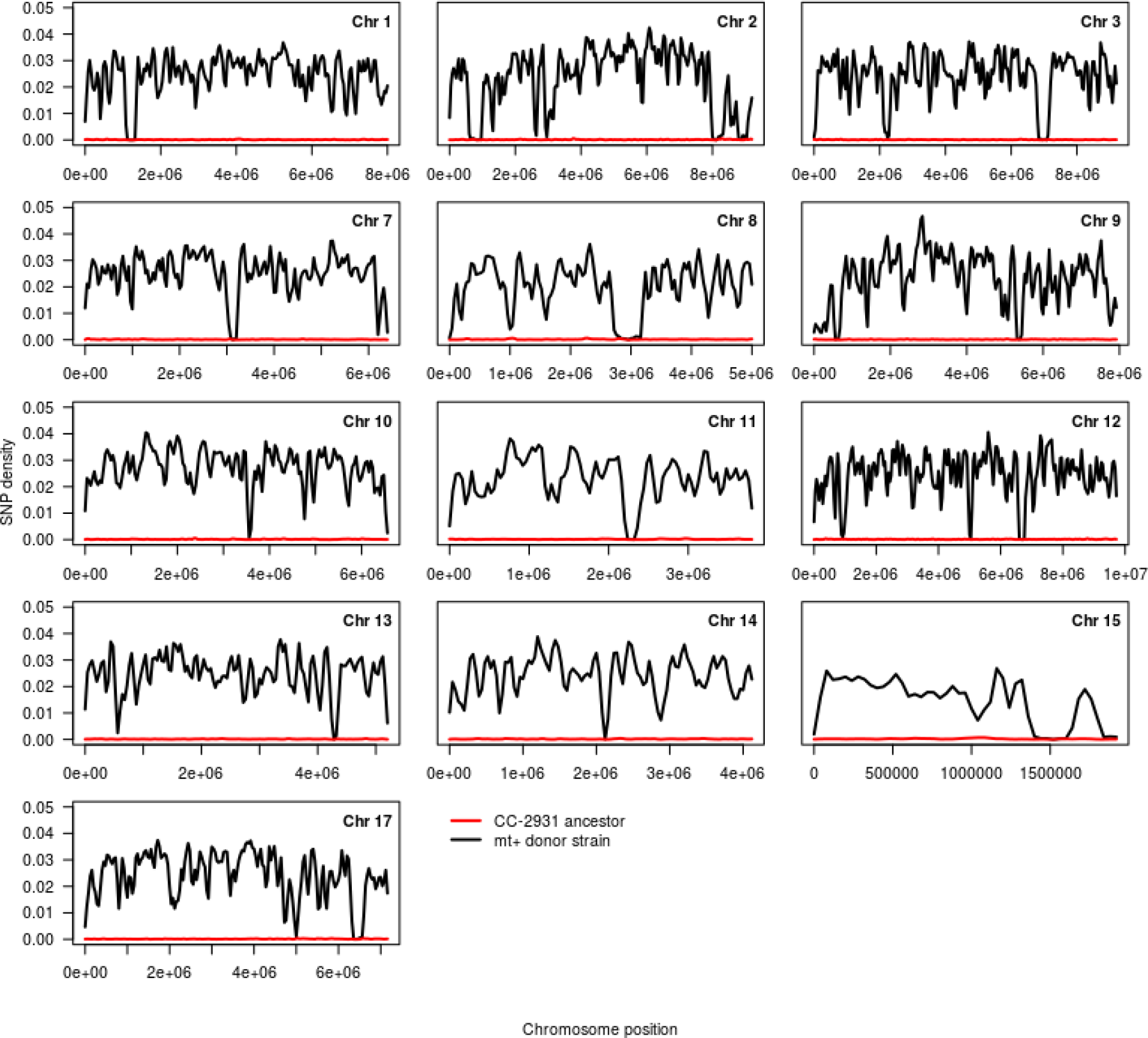
SNP densities along chromosomes 1 - 3, 7 - 15, and 17 between the compatible ancestor for CC-2931 and its two ancestral strains. SNP densities were calculated for 80 kb windows along the chromosomes between the compatible ancestor and CC-2931 (red) and between the compatible ancestor and the mating type + donor strain (black). A mutation density of 0 indicates no genetic differences between the compatible ancestor and the strain it was compared to.

**Figure S11.**
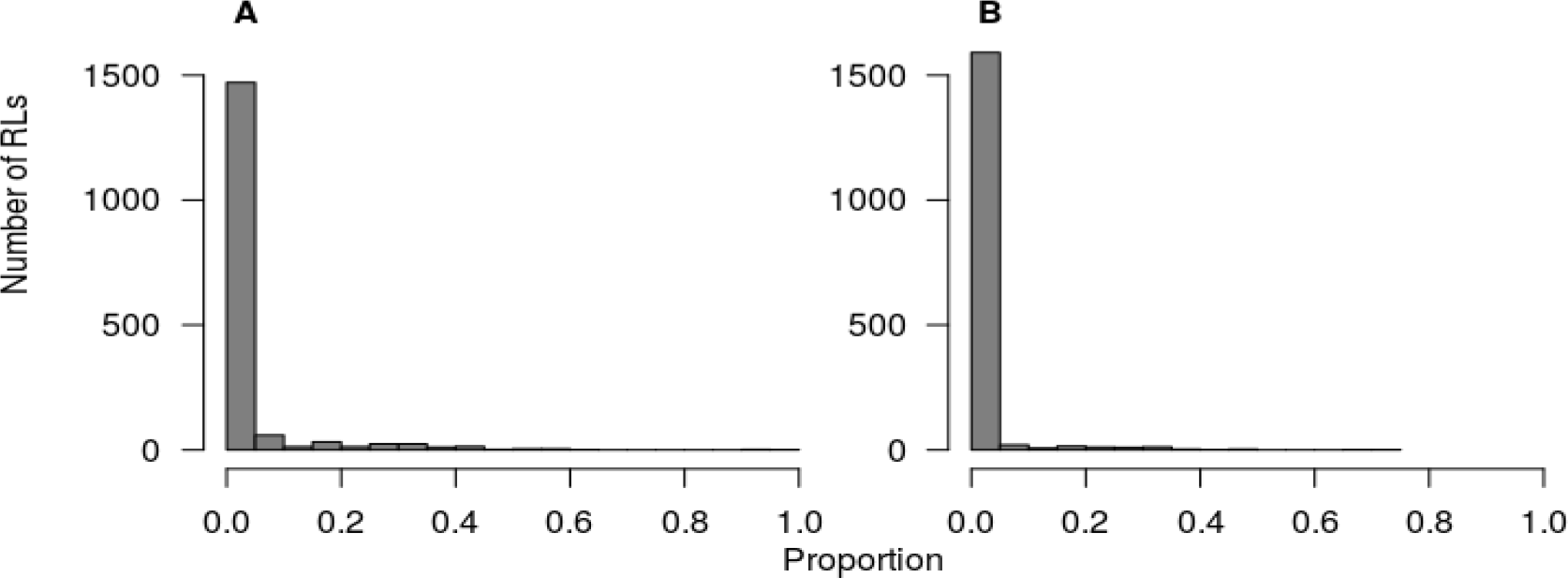
Distribution of missing data and heterozygous calls. The distribution of A) the proportion of missing data, i.e. non callable mutations across the whole data set, and of B) the proportion of heterozygous calls. Based on these distributions RLs with > 10% missing data and/or > 5% heterozygous calls were excluded from all analyses.

**Figure S12.**
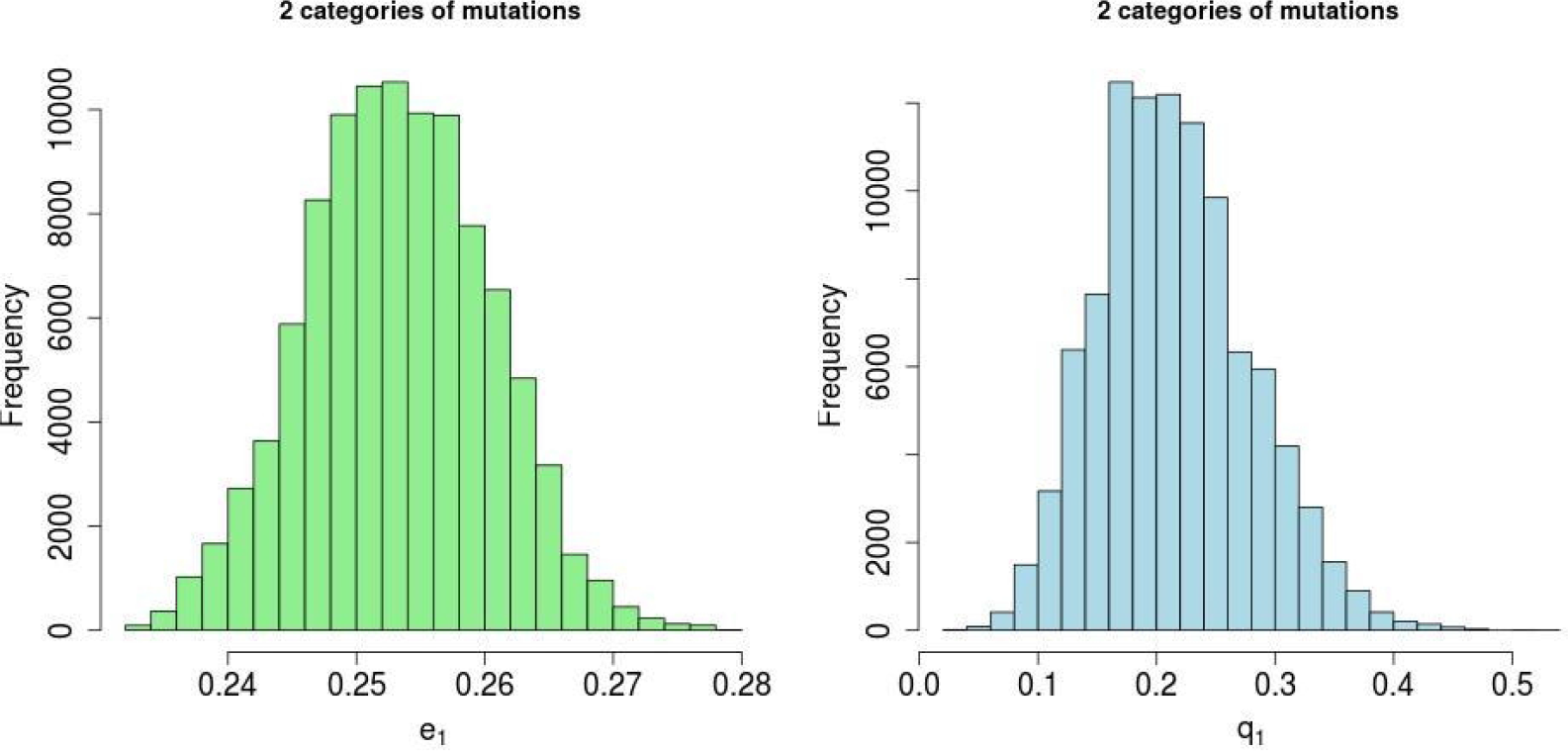
Posterior density plots for parameters *e*_1_ and *q*_1_ from MCMC analysis of simulated data with two categories of mutational effects, including one zero-effect category. The simulated values were *e*_1_ = 0.25 and *q*_1_ = 0.2. The mutation effect here and in Figs S2 and S3 are expressed in phenotypic standard deviation units. There were 40 mutations simulated and 10,000 observations.

**Figure S13.**
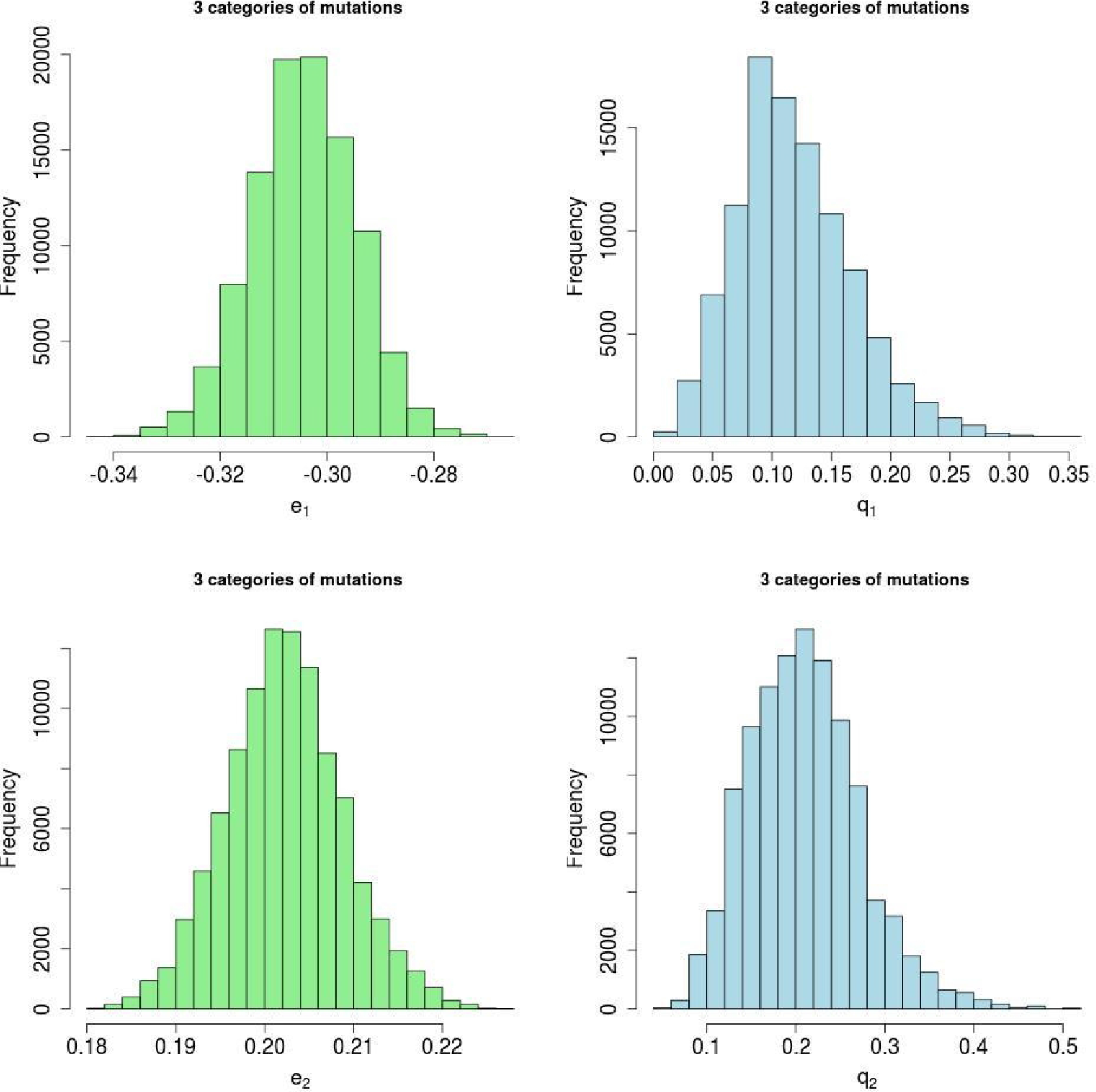
Posterior density plots for *e* and *q* parameters from MCMC analysis of simulated data with three categories of mutational effects, including one zero-effect category. The simulated values were *e*_1_ = −0.3, *q*_1_ = 0.1, *e*_2_ = 0.2 and *q*_2_ = 0.2.

**Figure S14.**
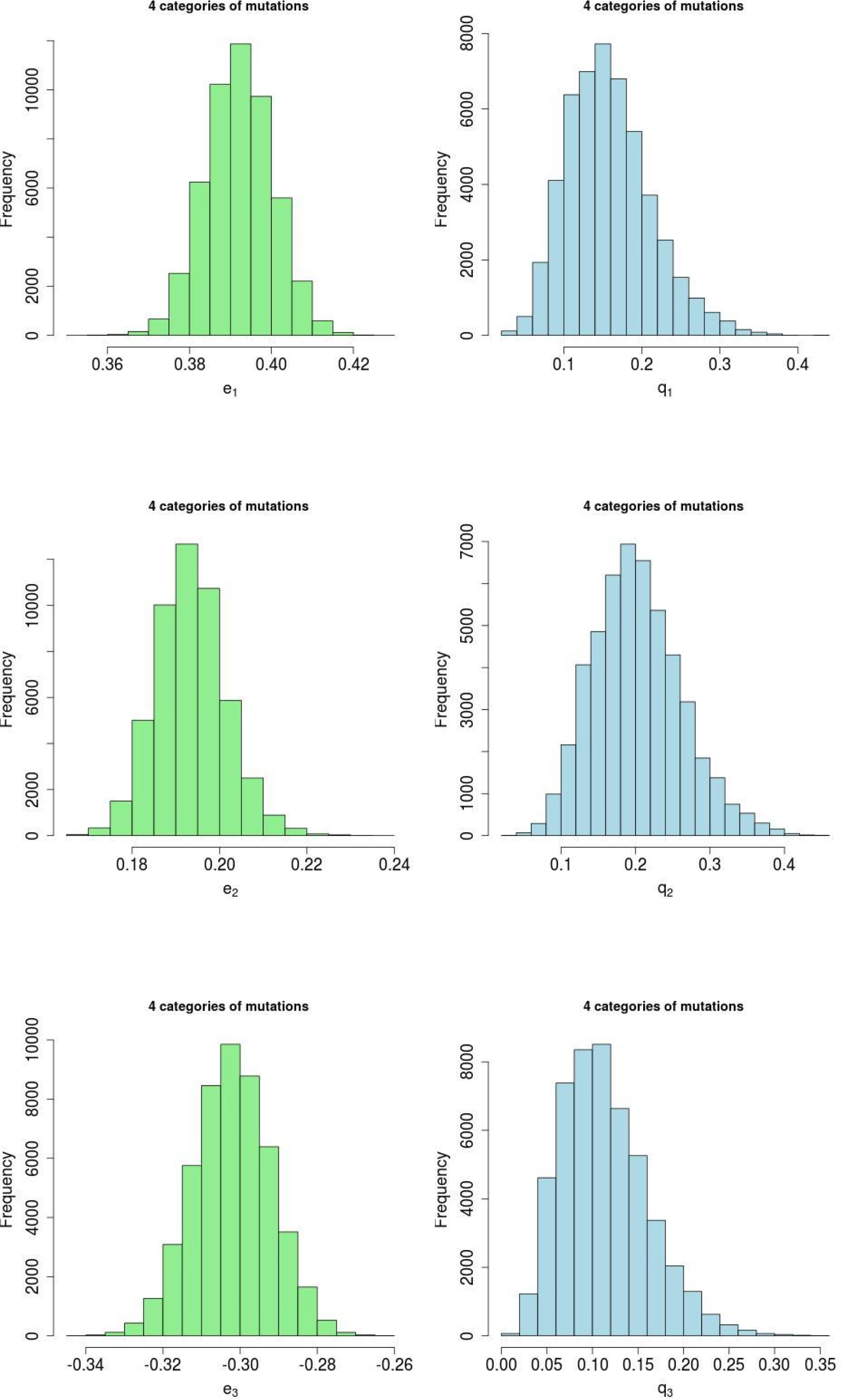
Posterior density plots for *e* and *q* parameters from MCMC analysis of simulated data with four categories of mutational effects, including one zero-effect category. The simulated values were *e*_1_ = 0.4, *q*_1_ = 0.15, *e*_2_ = 0.2, *q*_2_ = 0.2, *e*_3_ = −0.3 and *q*_3_ = 0.1.

**Figure S15.**
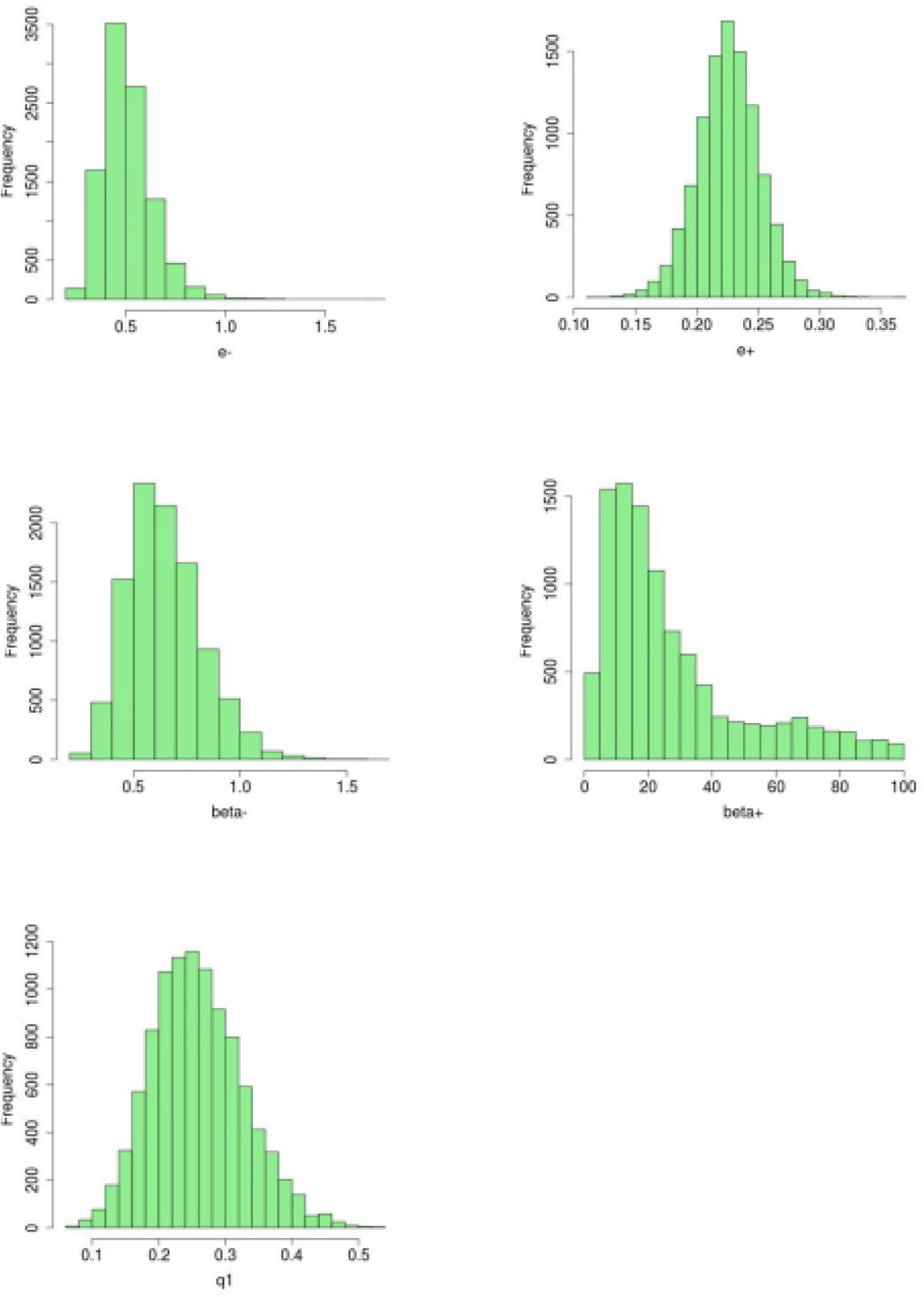
Posterior density plots for mean effects of negative (*e-*) and positive mutations (*e+*), gamma distribution shape parameters (*beta-* and *beta+) an*d the proportion of positive-effect mutations (*q1*) from MCMC analysis of simulated data under a two-sided gamma distribution of mutational effects. The simulated values were *e-* = 0.5, e+ = 0.25, beta- = 0.5, beta+ = 2, *q*_1_ = 0.25,.

### Supplementary tables

**Table S1.**
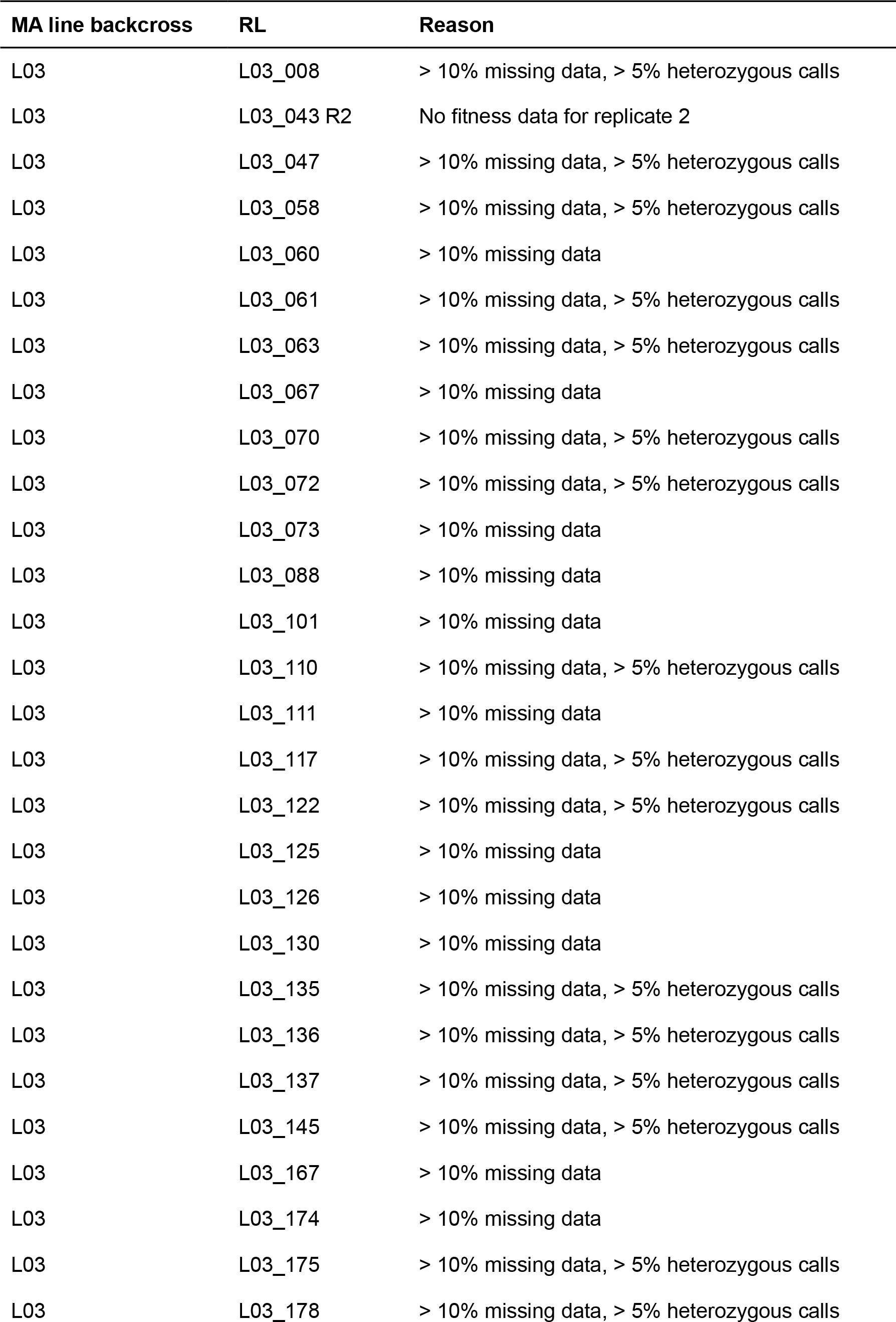
RLs that were excluded from all analyses.

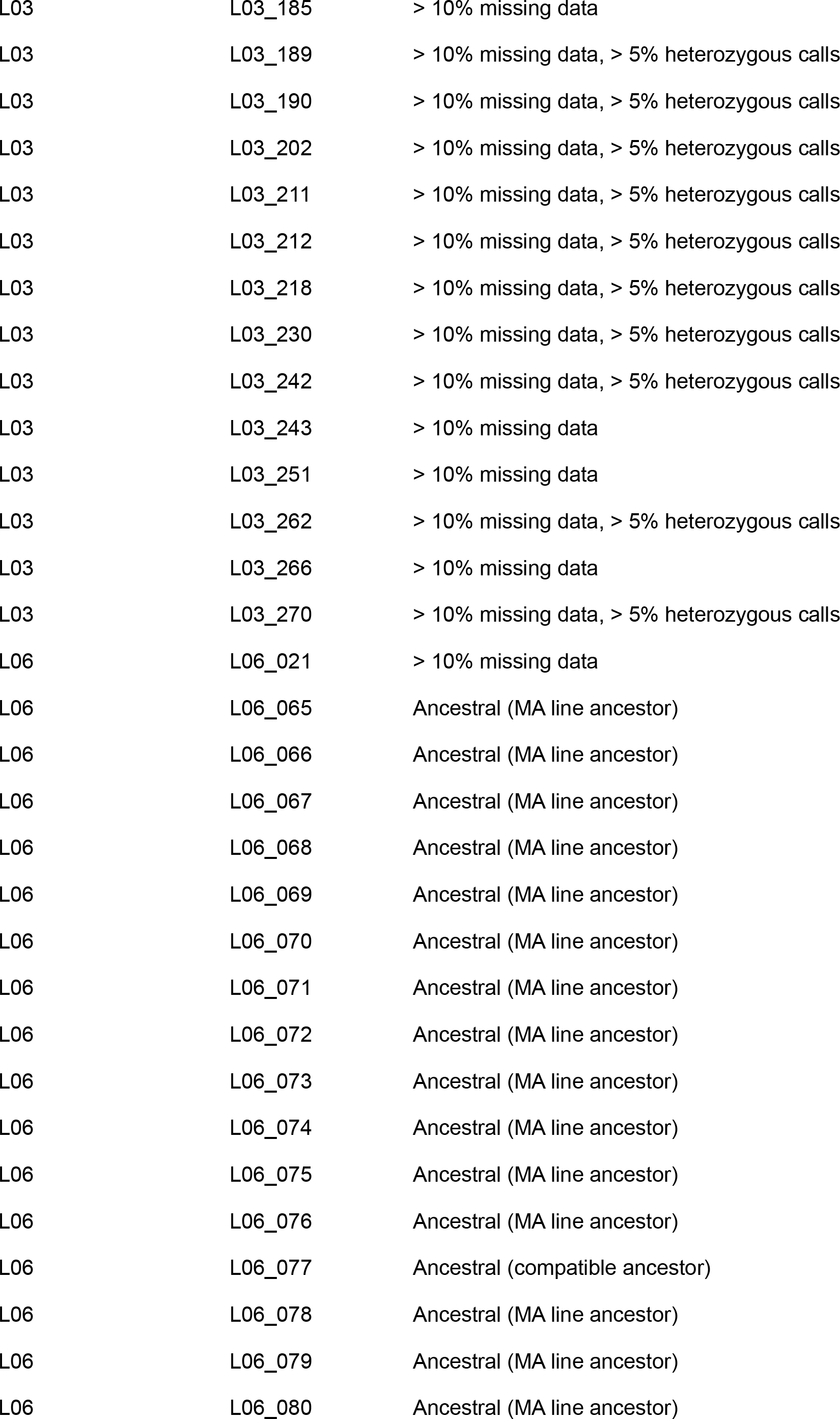

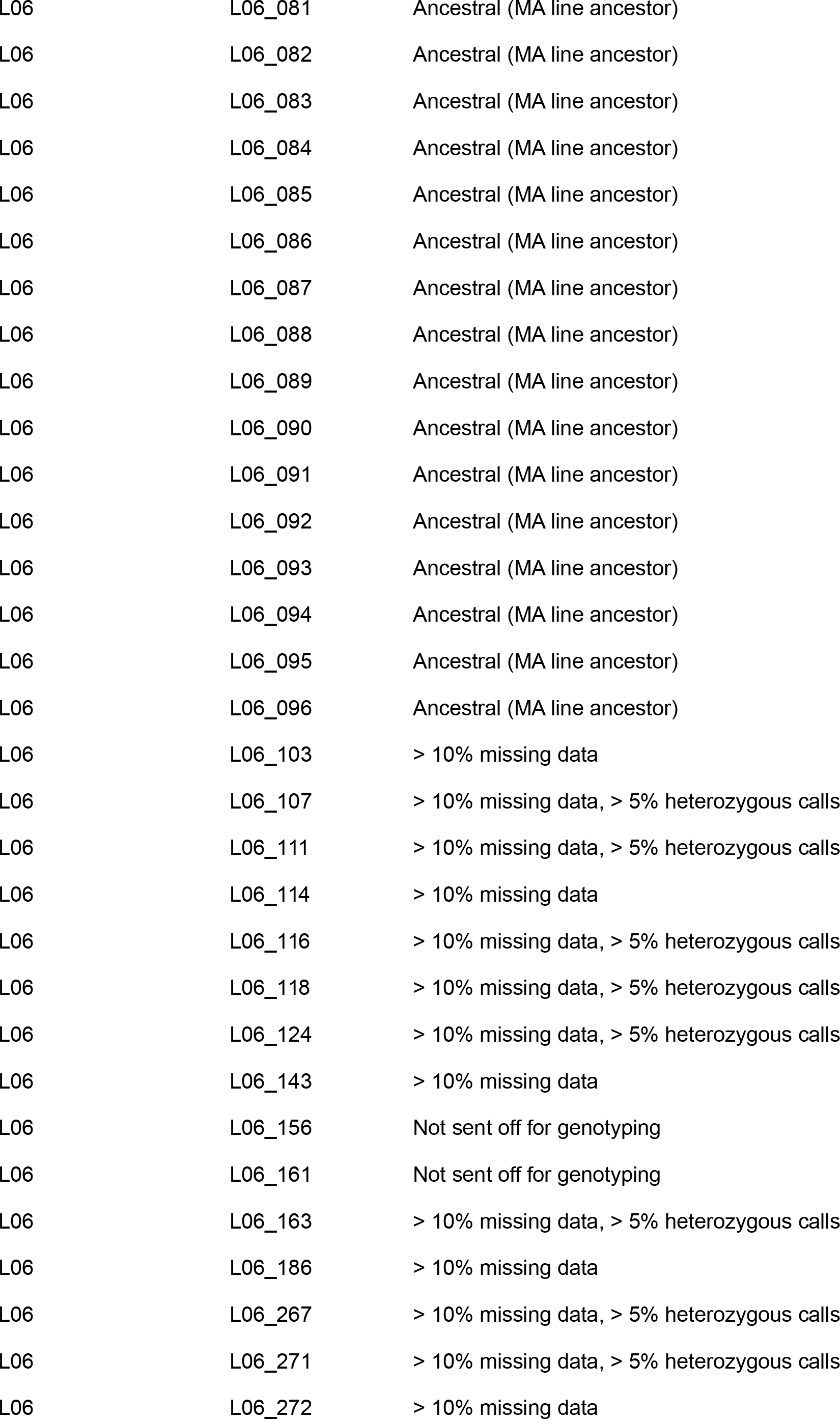

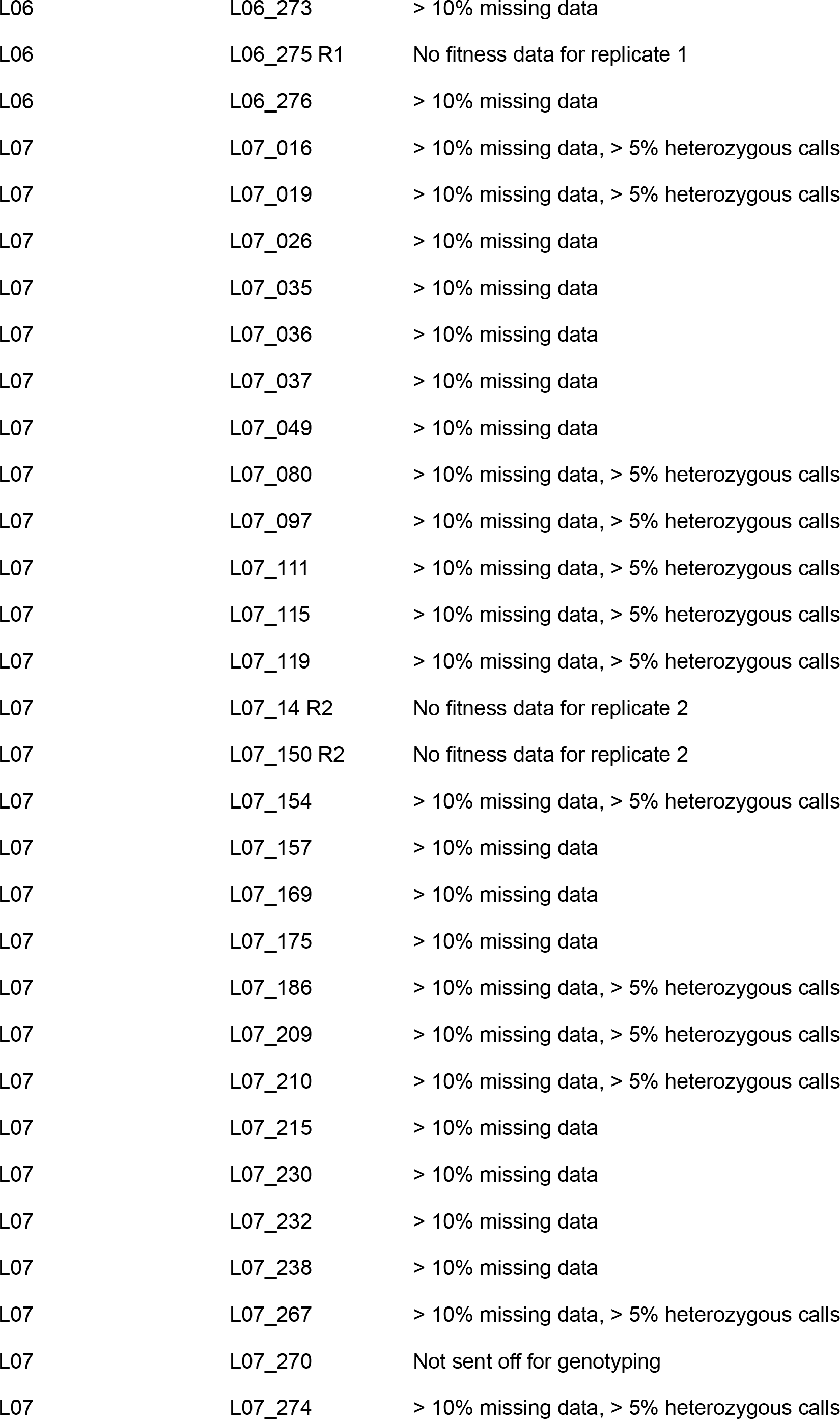

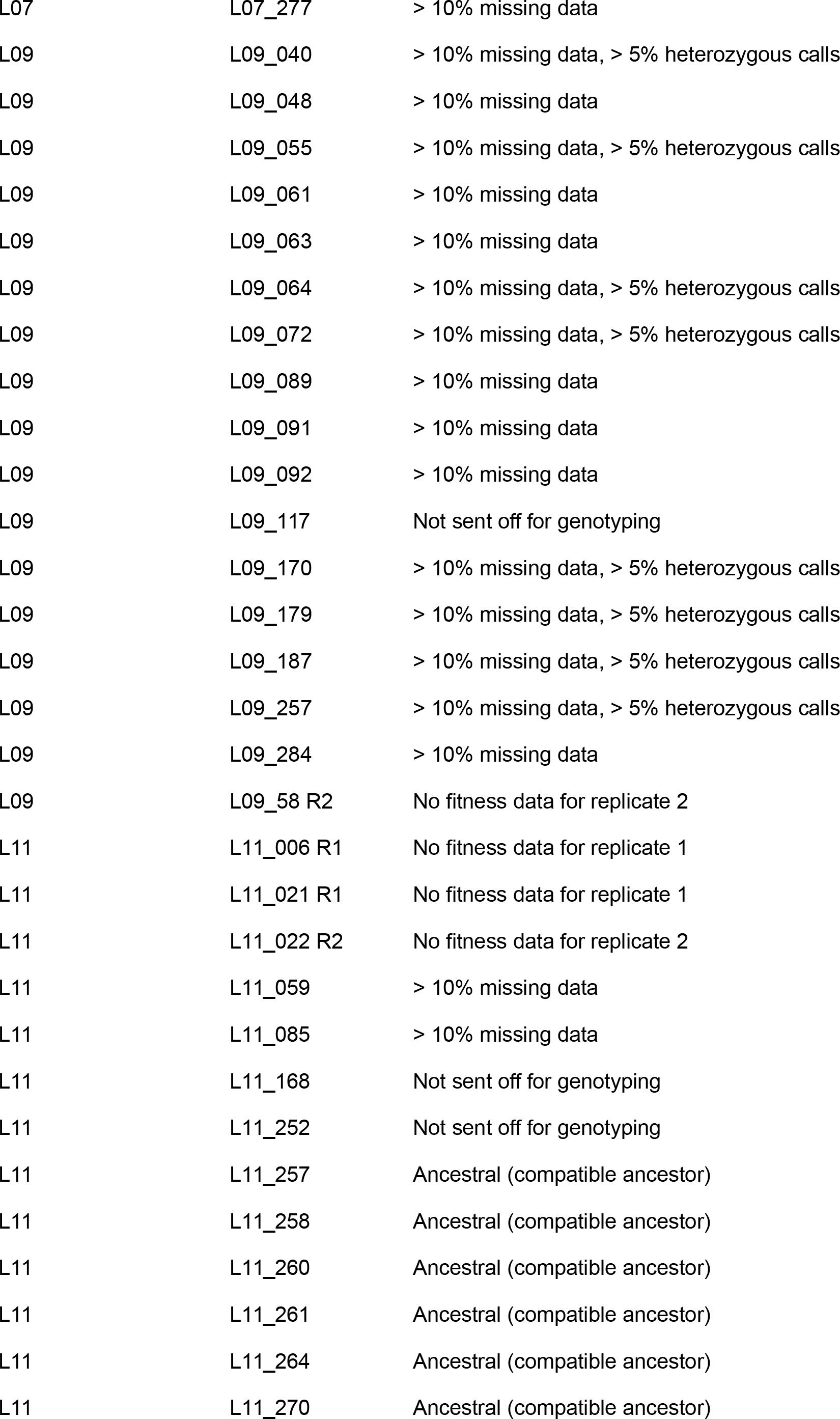

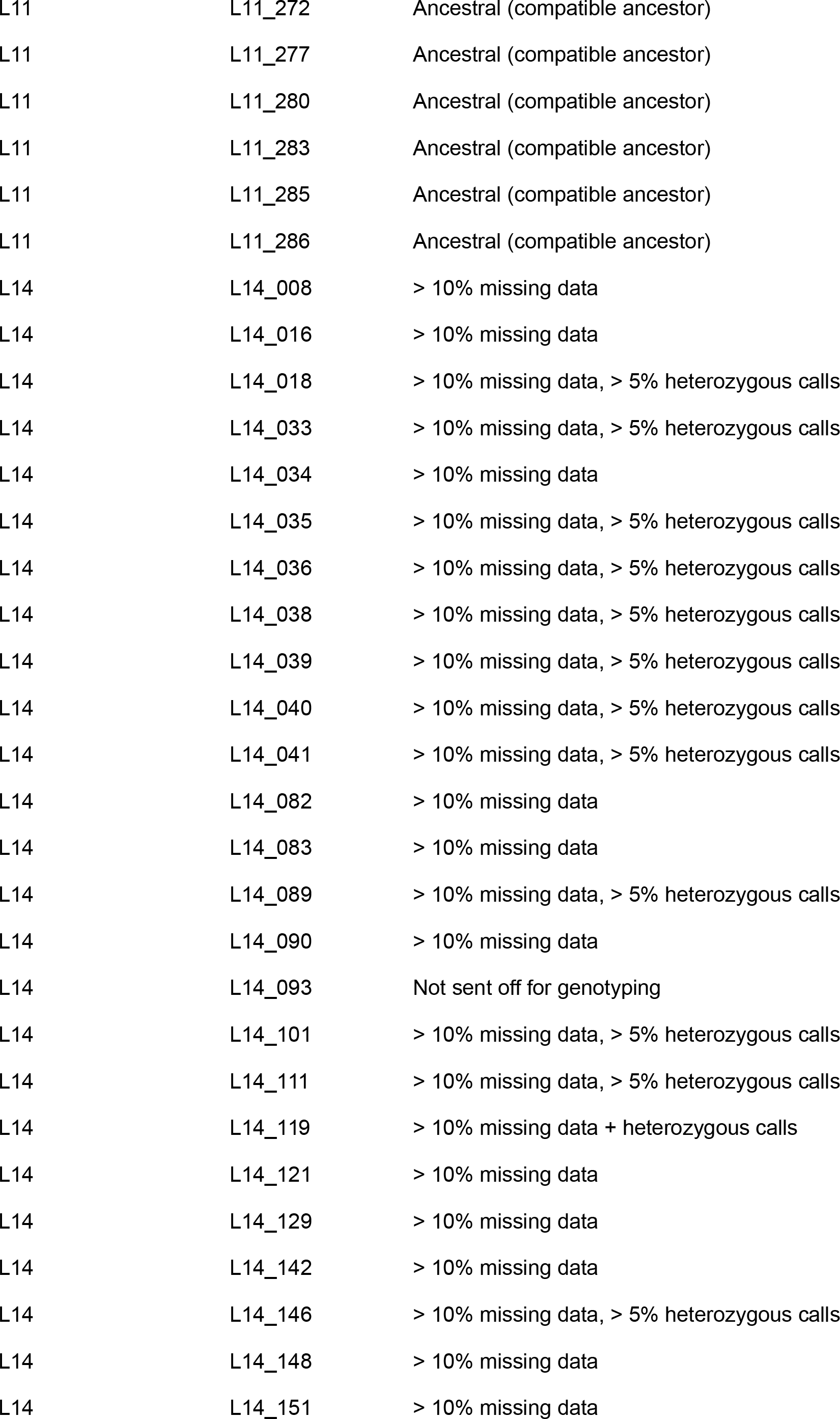

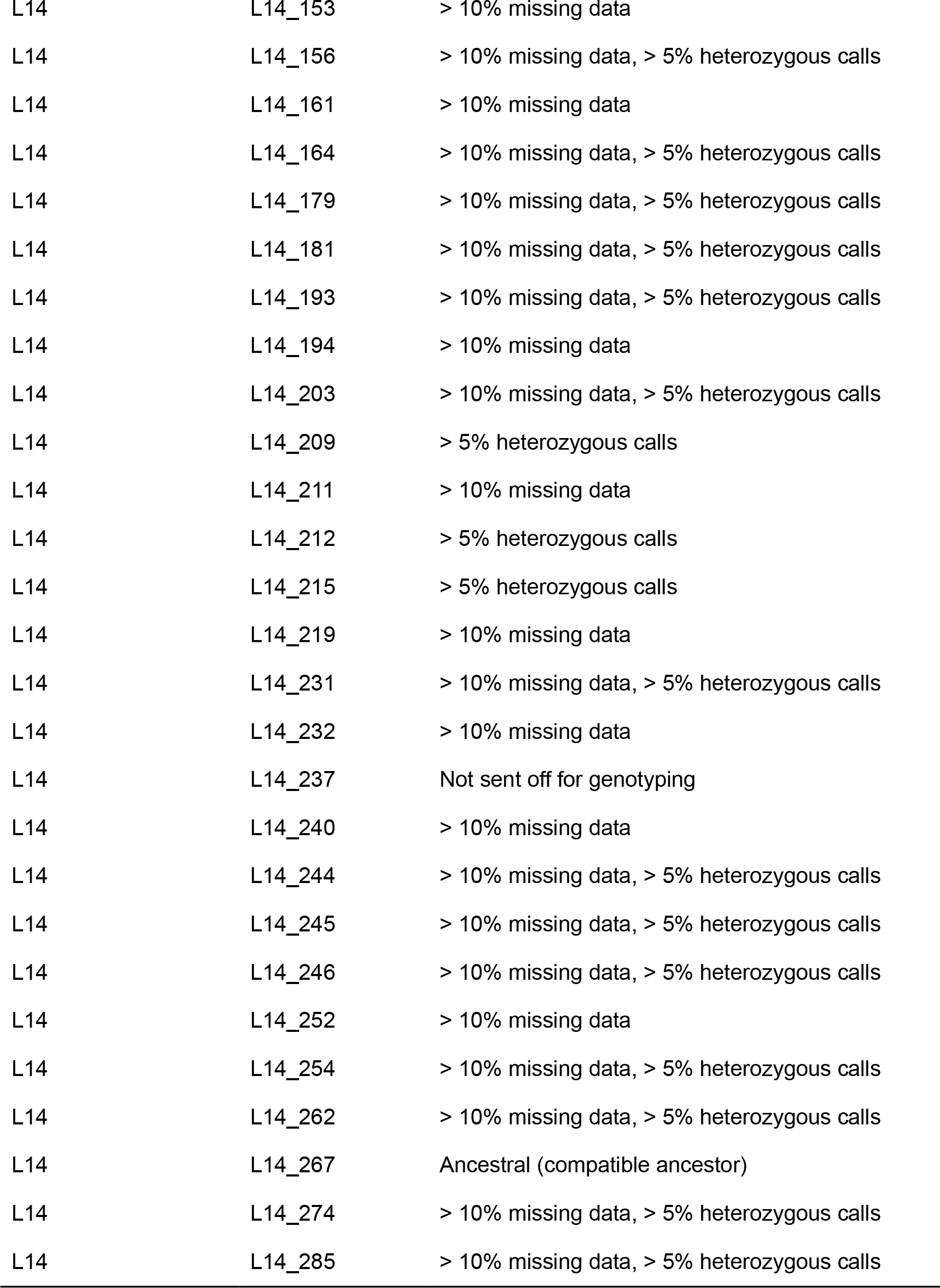

**Table S2.**
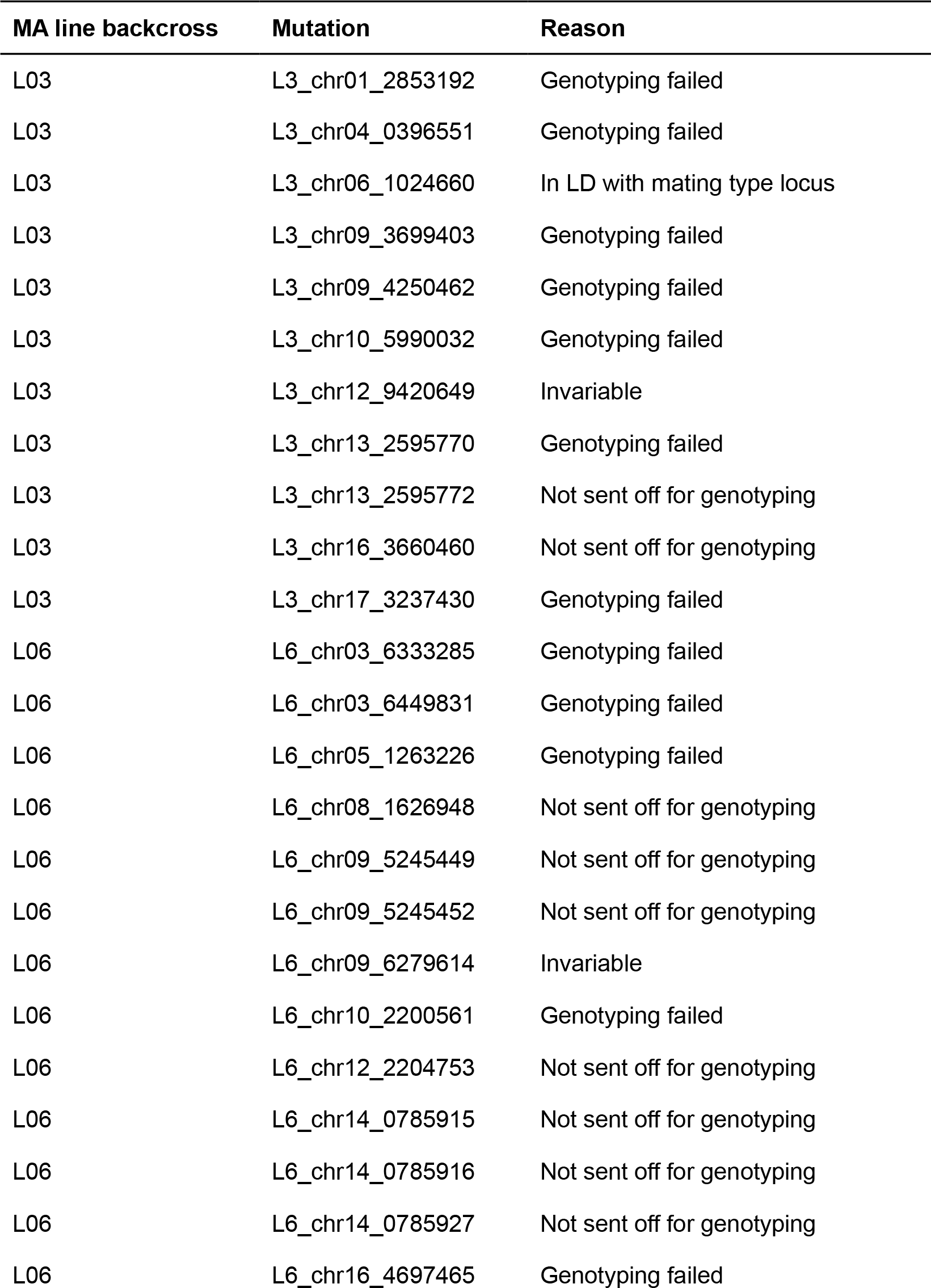
Mutations that were excluded from all analyses.

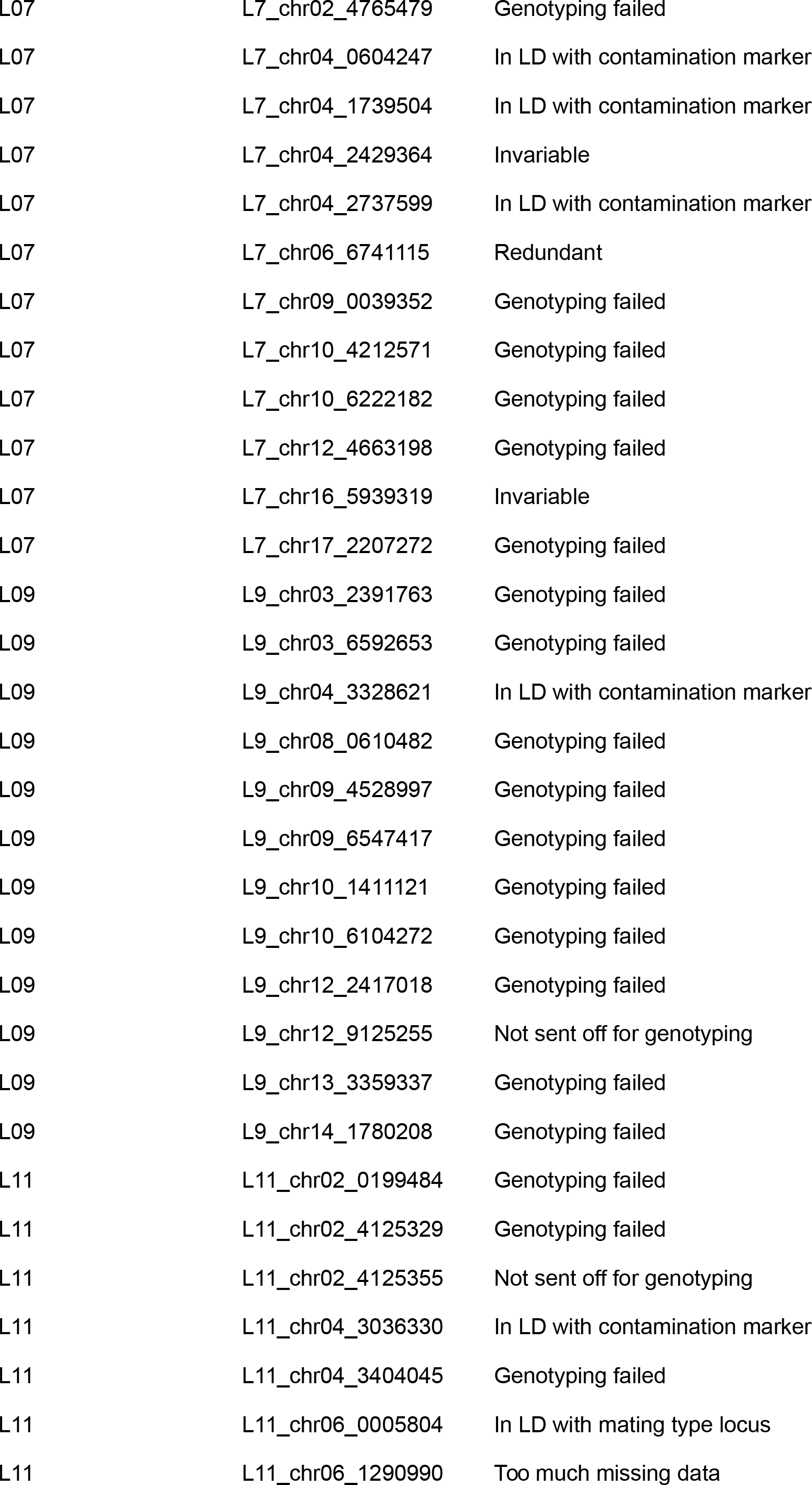

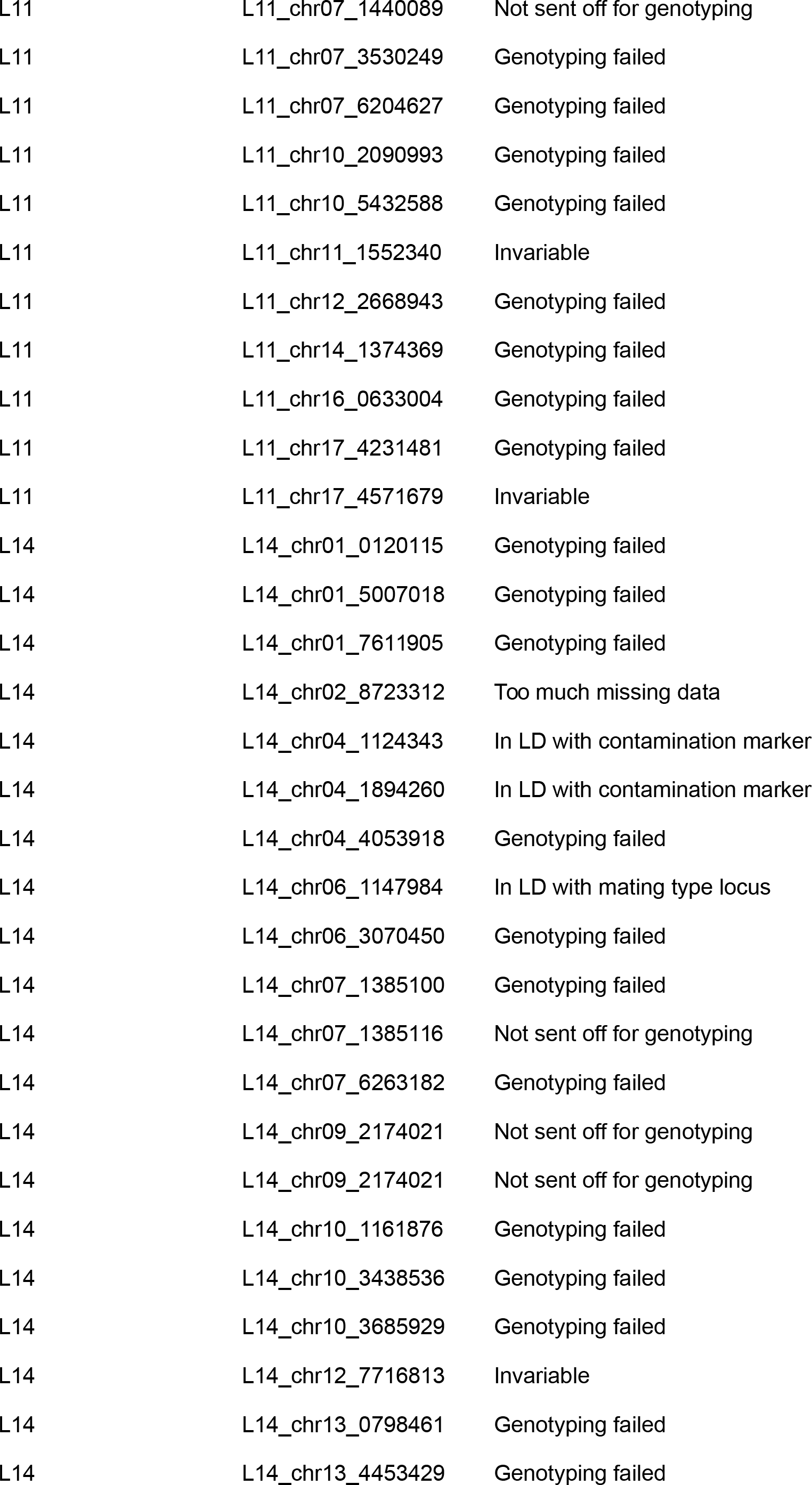

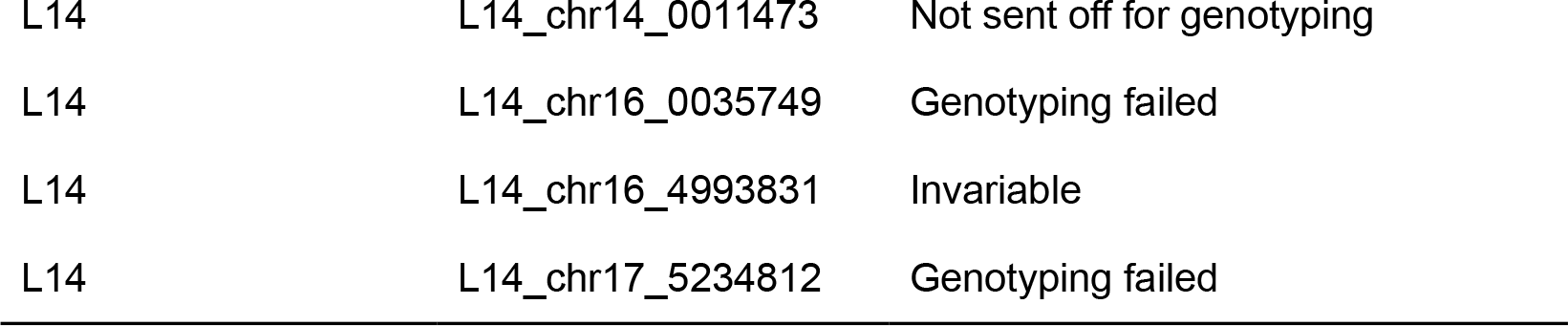

**Table S3.**
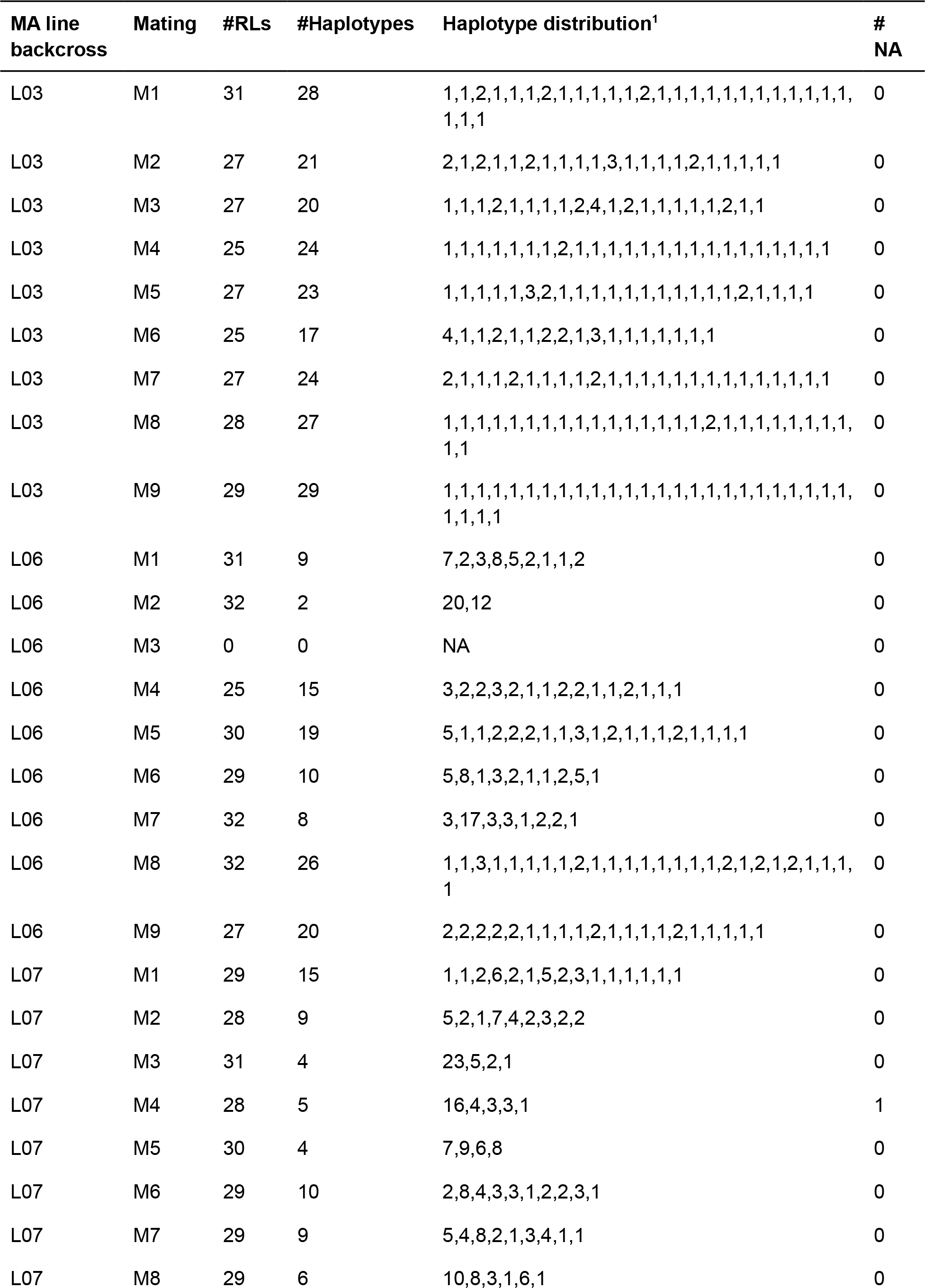
Number of haplotypes from each MA line backcross.

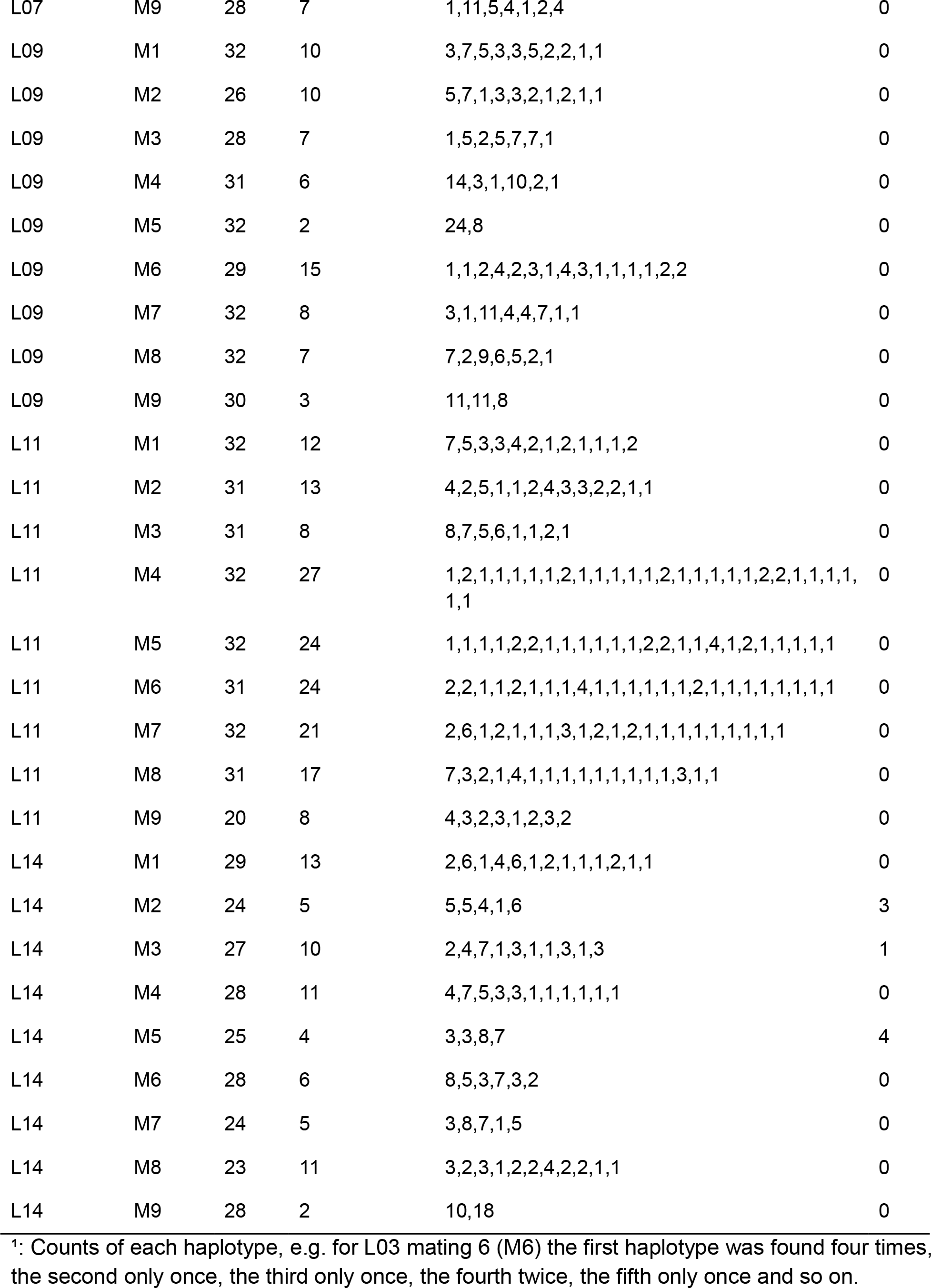

**Table S4.**
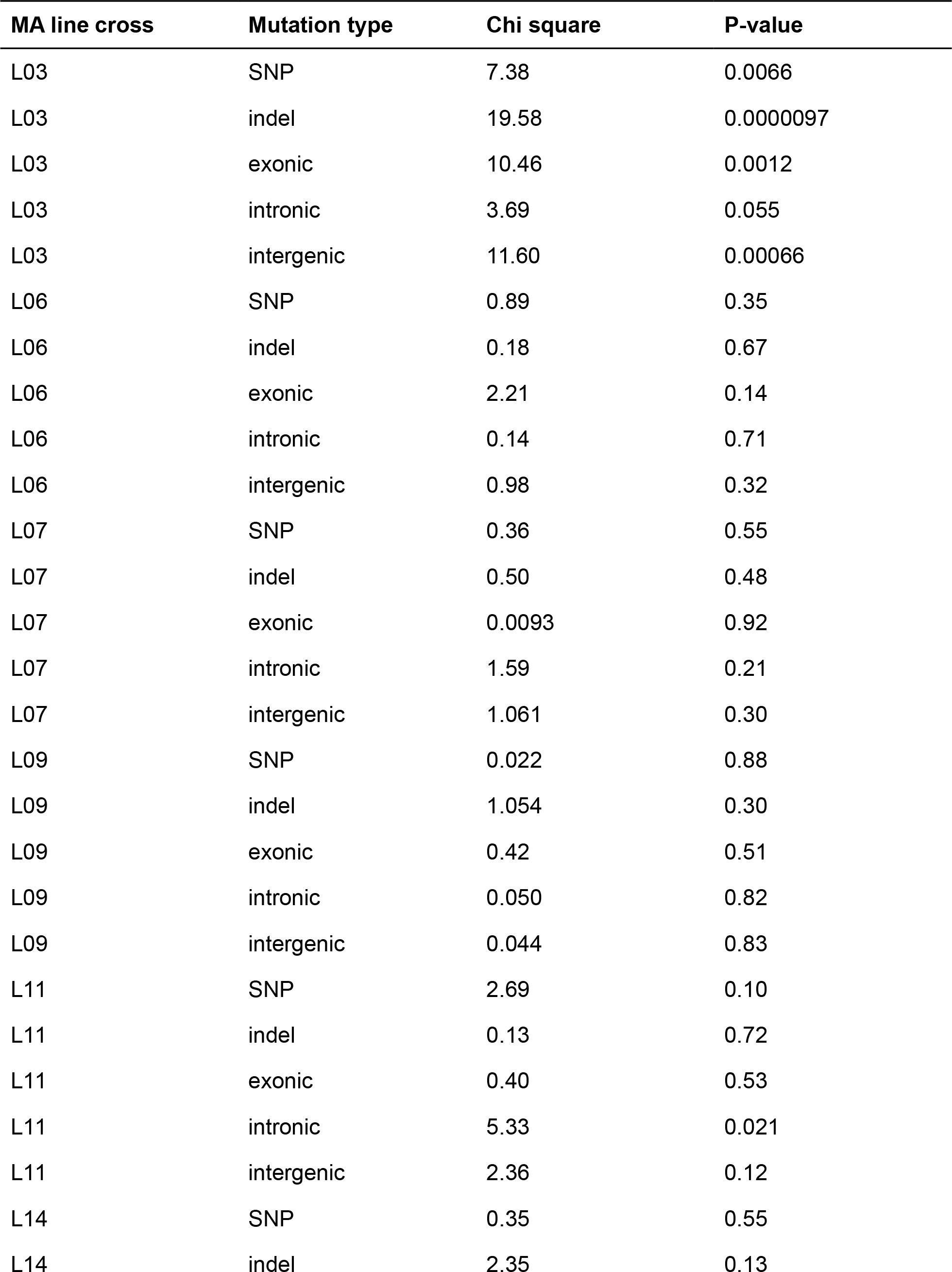
Likelihood ratio tests for mixed model analysis of growth rate as a function of number of different mutation types with 1 degree of freedom.

**Table S4.**
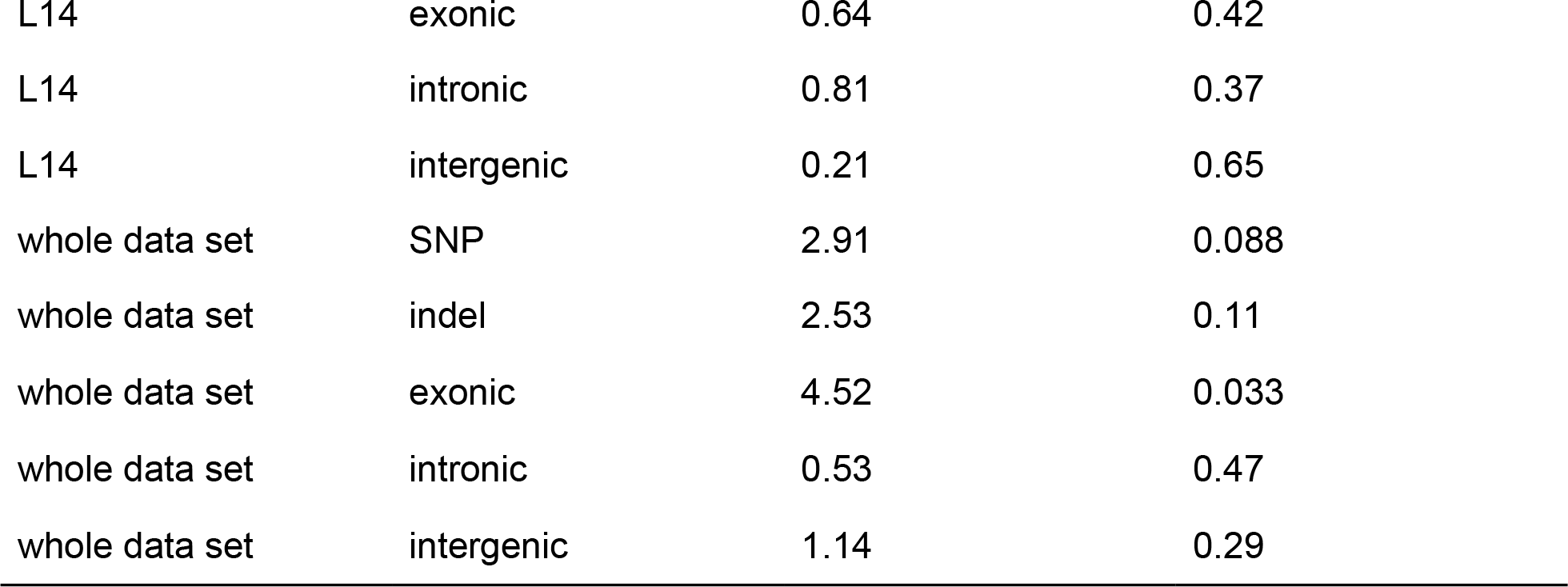

**Table S5.** Parameter estimates for the two mutation effect category model, including one zero-effect category.

| **MA Line** | ***Parameter estimate (95% credible interval)*** |  |
| --- | --- | --- |
|  | **$e_1$** | **$q_1$** |
| L03 | -0.018 (-0.040, -0.003) | 0.127 (0.031, 0.693) |
| L06 | 0.001 (-0.049, 0.060) | 0.017 (0.003, 0.729) |
| L07 | -0.019 (-0.063, 0.040) | 0.019 (0.005, 0.765) |
| L09 | 0.001 (-0.043, 0.044) | 0.002 (0.001, 0.819) |
| L11 | 0.051 (0.012, 0.072) | 0.032 (0.007, 0.181) |
| L14 | -0.038 (-0.061, -0.016) | 0.051 (0.016, 0.192) |

**Table S6.** (.csv) file (large-effects.csv) containing effects and mutation types of top 10 absolute effect mutations.

**Table S7.** Mutations that were corrected.

| **MA line backcross** | **Mutation** | **# ancestral calls** | **# derived calls** | **# missing data (NA)** | **Correction** |
| --- | --- | --- | --- | --- | --- |
| L03 | L3_chr05_3408017 | 0 | 150 | 138 | NA replaced with 0 |
| L03 | L3_chr09_4604654 | 0 | 139 | 149 | NA replaced with 0 |
| L03 | L3_chr11_0040103 | 0 | 142 | 146 | NA replaced with 0 |
| L03 | L3_chr12_7098772 | 177 | 0 | 111 | NA replaced with 1 |
| L06 | L6_chr05_2967344 | 132 | 0 | 156 | NA replaced with 1 |
| L06 | L6_chr10_4886371 | 0 | 110 | 178 | NA replaced with 0 |
| L06 | L6_chr10_5830415 | 120 | 0 | 168 | NA replaced with 1 |
| L06 | L6_chr15_0715589 | 138 | 0 | 150 | NA replaced with 1 |
| L09 | L9_chr05_0258561 | 0 | 164 | 124 | NA replaced with 0 |
| L09 | L9_chr05_2061109 | 0 | 156 | 132 | NA replaced with 0 |
| L09 | L9_chr05_3366061 | 0 | 201 | 87 | NA replaced with 0 |
| L09 | L9_chr06_8166934 | 167 | 0 | 121 | NA replaced with 1 |
| L09 | L9_chr09_3706134 | 0 | 158 | 130 | NA replaced with 0 |
| L09 | L9_chr11_3439747 | 153 | 0 | 135 | NA replaced with 1 |
| L09 | L9_chr17_4121513 | 0 | 110 | 178 | NA replaced with 0 |
| L09 | L9_chr17_4814616 | 159 | 0 | 129 | NA replaced with 1 |
| L11 | L11_chr02_0189093 | 0 | 138 | 150 | NA replaced with 0 |
| L11 | L11_chr02_1921472 | 155 | 0 | 133 | NA replaced with 1 |
| L11 | L11_chr08_0145992 | 134 | 0 | 154 | NA replaced with 1 |
| L11 | L11_chr17_0652393 | 134 | 0 | 154 | NA replaced with 0 |
| L14 | L14_chr04_0006715 | 121 | 0 | 167 | NA replaced with 1 |

## References

Bateman, A. J. (1959). The viability of near-normal irradiated chromosomes. Intern. J. Radiat. Biol. 1: 170–180.

Bates, D., Mächler, M., Bolker, B.M., Walker, S.C., (2015). Fitting Linear Mixed-Effects Models Using lme4. Journal of Statistical Software, 67(1), 1–48. doi:10.18637/jss.v067.i01

Bold, H.C., (1942). The cultivation of algae. The Botanical Review 8: 69–138

Boyko, A. R., Williamson, S. H., Indap, A. R., Degenhardt, J. D., Hernandez, R. D., Lohmueller, K. E., Adams, M. D., Schmidt, S., Sninsky, J. J., Sunyaev, S. R., White, T. J., Nielsen, R., Clark, A. G. and Bustamante, C. D. (2008). Assessing the evolutionary impact of amino acid mutations in the human genome. PLoS Genetics 4: e1000083.

Charlesworth B. and Charlesworth D. (2010). Elements of Evolutionary Genetics. Greenwood Village, Colorado: Roberts & Co.

Eyre-Walker, A., Woolfit, M. and Phelps, T. (2006) The distribution of fitness of new deleterious amino acid mutations in humans. Genetics 173: 891–900.

Eyre-Walker, A. and Keightley, P. D. (2007). The distribution of fitness effects of new mutations. Nature Reviews Genetics 8: 610–618.

Gallet, R., Cooper, T. F., Elena, S. F. and Lenormand, T. (2012). Measuring selection coefficients below 10−3: method, questions, and prospects. Genetics 190: 175–186.

García-Dorado, A. (1997). The rate and effects distribution of viability mutation in Drosophila: Minimum distance estimation. Evolution 51: 1130–1139.

Graur, D., Zheng, Y., Price, N., Azevedo, R. B. R., Zufall, R. A. and Elhaik, E. (2013). On the immortality of television sets: “function” in the human genome according to the evolution-free gospel of ENCODE. Genome Biol Evol 5: 578–590.

Halligan, D. L. and Keightley, P. D. (2009). Spontaneous mutation accumulation studies in evolutionary genetics. Annual Review of Ecology, Evolution and Systematics 40: 151–172.

Hill, W. G. (1982). Rates of change in quantitative traits from fixation of new mutations. Proc. Natl. Acad. Sci. USA 79: 142–145.

Jasra, A., Holmes, C. C. and Stephens, D. A. (2005). Markov Chain Monte Carlo methods and the label switching problem in Bayesian mixture modeling. Statist. Sci. 20: 50–67.

Katju, V. and Bergthorsson, U. (2018). Old trade, new tricks: insights into the spontaneous mutation process from the partnering of classical mutation accumulation experiments with high-throughput genomic approaches. Genome Biology and Evolution, evy252, https://doi.org/10.1093/gbe/evy252.

Keightley, P. D., (1994). The distribution of mutation effects on viability in Drosophila melanogaster. Genetics 138: 1315–1322.

Keightley, P. D. (1998). Inference of genome wide mutation rates and distributions of mutation effects for fitness traits: a simulation study. Genetics 150: 1283–1293.

Keightley, P. D. and Hill, W. G. (1990). Variation maintained in quantitative traits with mutation-selection balance: pleiotropic side-effects on fitness traits. Proceedings of the Royal Society of London Series B 242: 95–100.

Keightley, P. D. and Lynch, M. (2003). Towards a realistic model of mutations affecting fitness. Evolution 57: 683–685.

Keightley, P. D. and Eyre-Walker, A. (2007). Joint inference of the distribution of fitness effects of deleterious mutations and population demography based on nucleotide polymorphism frequencies. Genetics 177: 2251–2261.

Kimura, M. (1979). Model of effectively neutral mutations in which selective constraint is incorporated. Proc. Natl. Acad. Sci. USA 76: 3440–3444.

Kimura, M. (1983). The Neutral Theory of Molecular Evolution. Cambridge University Press.

Kraemer, S. A., Morgan, A. D., Ness, R. W., Keightley, P. D. and Colegrave, N. (2016). Fitness effects of new mutations in Chlamydomonas reinhardtii across two stress gradients. Journal of Evolutionary Biology 29: 583–593

Kraemer, S.A., Böndel, K.B., Ness, R.W., Keightley, P.D., Colegrave, N. (2017). Fitness change in relation to mutation number in spontaneous mutation accumulation lines of Chlamydomonas reinhardtii. Evolution 71: 2918–2929.

Leffler, E. M., Bullaughey, K., Matute, D. R., Meyer, W. K., Segurel, L., Venkat, A., Andolfatto, P. and Przeworski, M. (2012). Revisiting an old riddle: what determines genetic diversity levels within species? PLoS Biol 10: e1001388.

Merchant, S.S., Prochnik, S.E., Vallon, O., Harris, E.H., Karpowicz, S.J., Witman, G.B., Terry, A., Salamov, A., Fritz-Laylin, L.K., Maréchal-Drouard, L., et al. (2007). The Chlamydomonas genome reveals the evolution of key animal and plant functions. Science 318: 245–250.

Morgan, A.D., Ness, R.W., Keightley, P.D., Colegrave, N. (2014). Spontaneous mutation accumulation in multiple strains of the green alga, Chlamydomonas reinhardtii. Evolution 68: 2589–2602.

Moser, G, Lee, SH, Hayes, BJ, Goddard, ME, Wray, NR and Visscher PM (2015) Simultaneous discovery, estimation and prediction analysis of complex traits using a Bayesian mixture model. PLoS Genet 11: e1004969.

Mukai, T. (1964). The genetic structure of natural populations of Drosophila melanogaster. I. Spontaneous mutation rate of polygenes controlling viability. Genetics 50: 1–19.

Ness, R.W., Morgan, A.D., Vasanthakrishnan, R.B., Colegrave, N., Keightley, P.D., (2015). Extensive de novo mutation rate variation between individuals and across the genome of Chlamydomonas reinhardtii. Genome Research 25: 1739–1749.

Ohta, T. (1977) Extension of the neutral mutation drift hypothesis. In: Kimura M (ed) Molecular Evolution and Polymorphism. National Institute of Genetics, Mishima, pp. 148–167.

Postma, E. (2014). Four decades of estimating heritabilities in wild vertebrate populations: Improved methods, more data, better estimates? Quantitative Genetics in the Wild. A. Charmantier, D. Garant and L. E. B. Kruuk. Oxford, Oxford University Press: p16–33.

R Core Team (2018). R: A language and environment for statistical computing. R Foundation for Statistical Computing, Vienna, Austria. URL https://www.R-project.org/.

Robert, L., Ollion, J., Robert, J., Song, X. H., Matic, I. and Elez, M. (2018). Mutation dynamics and fitness effects followed in single cells. Science 359: 1283–1286.

Robertson, A. (1967). The nature of quantitative genetic variation. pp. 265–280. In: Heritage from Mendel. (Ed. R. B. Brink). Madison, Milwaukee and London: University of Wisconsin Press.

Rutter, M. T., Roles, A. J. and Fenster, C. B. 2018 Quantifying natural seasonal variation in mutation parameters with mutation accumulation lines. Ecol Evol. 8: 5575–5585.

Sager, R. and Granick, S. (1954). Nutritional control of sexuality in Chlamydomonas reinhardtii. The Journal of General Physiology 37: 729–742.

Sanjuan, R., Moya, A. and Elena, S. F. (2004). The distribution of fitness effects caused by single-nucleotide substitutions in an RNA virus. Proc. Natl Acad. Sci. USA 101, 8396–8401.

Schneider, A., Charlesworth, B., Eyre-Walker, A. and Keightley, P. D. (2011). A method for inferring the rate of occurrence and fitness effects of advantageous mutations. Genetics 189: 1427–1437.

Shaw, F. H., Geyer C. J. and Shaw, R. G. (2002). A comprehensive model of mutations affecting fitness and inferences for Arabidopsis thaliana. Evolution 56: 453–463.

Tataru, P., Mollion, M., Glémin, S. and Bataillon, T. (2017). Inference of distribution of fitness effects and proportion of adaptive substitutions from polymorphism data. Genetics 207: 1103–1119.

